# DORSAL RAPHE NUCLEUS CONTROLS MOTIVATIONAL STATE TRANSITIONS IN MONKEYS

**DOI:** 10.1101/2024.02.13.580224

**Authors:** Luke Priestley, Mark Chiew, Mo Shahdloo, Ali Mahmoodi, Xinghao Cheng, Robin Cleveland, Matthew Rushworth, Nima Khalighinejad

## Abstract

The dorsal raphe nucleus (DRN) is an important source of serotonin in the brain but fundamental aspects of its function remain elusive. Here, we present a combination of minimally invasive recording and disruption studies to show that DRN brings about changes in motivation states. We use recently developed methods for identifying temporal patterns in behaviour to show that monkeys change their motivation depending on the availability of rewards in the environment. Distinctive patterns of DRN activity occur when monkeys transition between a high motivation state occupied when rewards are abundant, to a low motivation state engendered by reward scarcity. Disrupting DRN diminishes sensitivity to the reward environment and perturbs transitions in motivational states.

## INTRODUCTION

Animals need rewards for their survival, and they need to obtain them as efficiently as possible. In many naturalistic scenarios, this means tracking general features of the surrounding environment: foraging behaviour in many species, for example, involves comparing the opportunity an animal is currently confronted with against the general richness and stochasticity of opportunities it has encountered in the recent past, which guide its expectations for the future^1,2^. These are rational considerations for animals given the biological constraints that encumber them. Finding and pursuing rewards consumes precious metabolic resources that must later be replenished, and so it is critical that animals organise their reward-seeking activities in ways that exploit their external milieu – to press their advantage when things are good, and conserve their energy when things are bad.

How does the brain reconcile motivation for rewards with the environment in this way? Here, we argue for a critical role for the dorsal raphe nucleus (DRN) – a phylogenetically ancient part of the brainstem that is distinguished by its serotonergic innervation of the mammalian forebrain^3–6^. Although fundamental aspects of DRN’s function remain elusive, recordings of its activity during decision-making are beginning to cohere into two major themes: (i) that DRN controls changes in an animal’s behaviour^7–14^, and (ii) that DRN responds to reward-related features of an animal’s milieu, like the value, valence and uncertainty of recent outcomes^15–23^.

We build on previous work in arguing that DRN controls transitions between motivational states that reconcile an animal’s behaviour with the distribution of rewards in its environment. We develop a novel behavioural paradigm which demonstrates that rhesus monkeys are more motivated to pursue rewards that occur in rich environments with many high-value opportunities. We implement recent innovations in quantitative modelling to show that this pattern is explicable by state-like changes in an animal’s motivation that are consonant with the current distribution of rewards in the environment. We take advantage of the whole-brain perspective afforded by functional magnetic resonance imaging (fMRI) to show that brain activity in DRN – but no other neuromodulatory nucleus – covaries with transitions in motivation-states as well as the aspects of the environment on which they depend. Finally, we show that DRN is causally involved in motivational-state transitions by modulating neural activity with minimally invasive transcranial ultrasound stimulation^24,25^. In doing so, we provide the first demonstration that minimally invasive modulation of DRN is possible, and a new perspective on DRN’s behavioural function.

## RESULTS

### Motivation for rewards is modulated by the environment

Four rhesus monkeys (*Macaca mulatta*) performed a simple decision-making task involving sequential encounters with reward opportunities that varied in reward-magnitude and reward-probability (fig. 1A). Upon each encounter animals could either pursue the opportunity and incur a short temporal cost or let the opportunity pass and proceed to the next encounter. Each session comprised four blocks of 40-50 trials. We systematically controlled the distribution of reward-probabilities within blocks to engender different reward environments. These environments varied on two dimensions: (1) *richness*, defined by the mean reward-probability of opportunities– i.e., the value of an average opportunity in a block, and (2) *stochasticity*, defined as trial-to-trial variability in reward-probability, and implemented by changing the width of reward-probability distributions (fig. 1B & 1C; see Methods for details).

**Figure 1.**
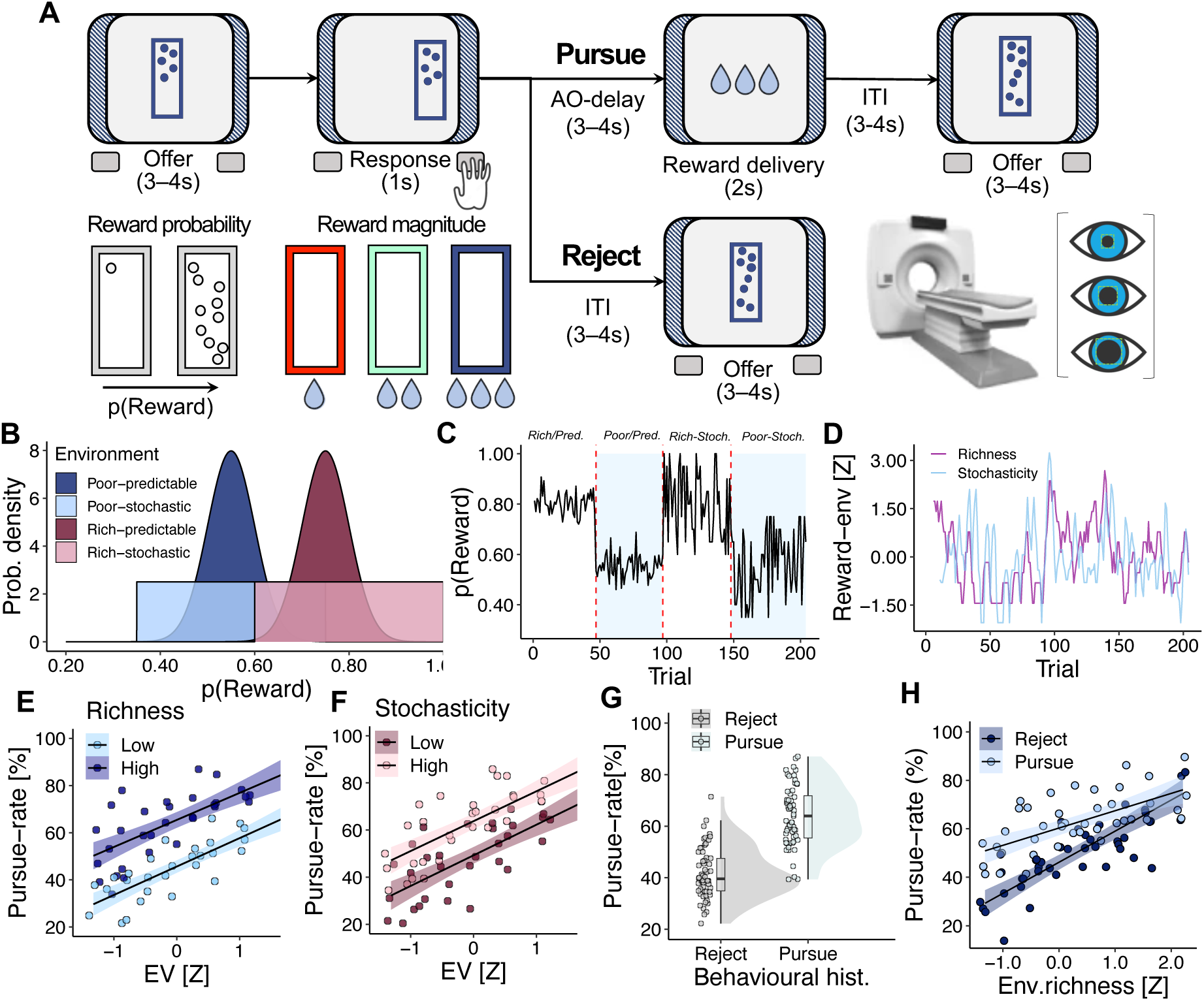
Animals are more likely to pursue reward opportunities in rich environments. **(A)** Animals performed a behavioural task involving sequential encounters with reward opportunities that varied in reward-magnitude (stimulus-colour) and reward-probability (dots-per-stimulus). Animals performed the task while undergoing functional magnetic resonance imaging and pupillometry **(B)** The probability density of reward-probability in different reward environments. Rich environments (purple; mean(reward-probability)=.75) were characterised by higher mean reward-probability than poor environments (blue; mean(reward-probability)=.55). Stochastic environments (light colours; sd(reward-probability)=.13) were characterised by uniform distributions and predictable environments by normal distributions (dark colours; sd(reward-probability)=.05). **(C)** A depiction of changes in reward-probability as a function of reward environment in an example session. Each session comprised four environments covering each permutation of richness and stochasticity. **(D)** Illustration of levels of the key dimensions for the example in previous panel: (i) richness and (ii) stochasticity. **(E)** Animals were more likely to pursue opportunities as the richness of the environment increased. X-axis indicates the expected value of the reward opportunity on each trial. Y-axis indicates rate of reward-pursuit. Colour-scale indicates mean-split according to richness of the environment. i.e. high corresponds to trials where richnesst > μrichness and low to richnesst ≤ μrichness. Dots indicate mean pursue-rates in quintile bins of expected-value. (**F)** Animals were more likely to pursue opportunities as the stochasticity of the environment increased. X-axis, Y-axis, dots and colour-scale follow conventions of **(E)**. **(G)** Animals were likely to repeat their previous decision over consecutive trials. Dots indicate mean level of responding per animal per session. In boxcharts, box-lengths indicate interquartile (IQR) ranges, bold lines indicate medians, and whiskers indicate median ± 1.5xIQR. **(H)** Behavioural history modulated the effect of richness of the environment. Dots indicate mean level of responding in quintiles of richness, split according to behavioural history.

We first examined whether animals changed their behaviour in response to the environment. We reasoned that knowledge of the environment would derive from an animal’s reward history and therefore operationalised the environment’s richness as the average reward accumulated in the previous five trials and its stochasticity as the standard deviation of the reward rate (fig. 1D). We selected a five-trial window because each individual animal showed sensitivity to individual reward outcomes up to five trials into the past (see supplementary figure S1B). We quantified the impact of reward environments on motivation with a series of binomial general linear models (GLM) in which motivation for rewards was conceived as the binomial likelihood of pursuing a reward opportunity. This revealed that animals were more likely to pursue opportunities as the environment’s richness and stochasticity increased (GLM1.1, see Methods; β_environment-richness_=0.46, SE=0.08, *p*<.001; fig.1E; see also supplementary figure S1A & S1C; β_environment-stochasticity_=0.08, SE=0.03, *p*=.020; fig.1F; see also supplementary figure S1D). These effects obtained independently of trial-by-trial reward value – for example, opportunities with the same expectation-value were more likely to be pursued in rich relative to poor environments, meaning that the environment effect was not an artefact of specific reward opportunity values on the current trial (fig.1E & 1F; see also supplementary figure S1A & C–D).

This analysis gestured toward a subtle but important feature of behaviour: animals could only obtain rewards by pursue-responses, and so the association between receiving rewards in the past and pursuing rewards in the present implied that behaviour was autocorrelated. We quantified this by regressing the pursue/reject decision taken on each trial against the corresponding decision on the previous trial (behavioural history, hence). This confirmed that animals were apt to repeat pursue/reject decisions over consecutive trials, despite trials featuring separate reward opportunities with distinct magnitude and probability parameters (GLM1.2; β_behavioural-history_=0.90, SE=0.24, *p*<.001; fig.1G). Although behavioural history and richness of the environment were correlated (*r*=0.43), they constituted separate and distinctive effects on behaviour according to a binomial GLM featuring both predictors (GLM1.3; β_richness_=0.54, SE=0.08, *p*<.001; β_behavioural-history_=0.58 SE=0.21, *p*=.005). This GLM revealed, moreover, an interaction whereby behavioural history’s influence diminished as the richness of the environment increased (GLM1.3; β=-0.33, SE=0.11, *p* =.002; fig.1H). This suggests that animals experienced time-varying predispositions to either to pursue or reject opportunities, and that their tendency to behave in either manner depended on the distribution of rewards in the recent past.

### A Hidden Markov Model identifies motivation-states in behaviour

We reasoned that these behavioural patterns reflected changes in internal motivation-states – that is, changes in an animal’s intrinsic propensity to pursue rewards independent of the value of the opportunity currently at stake (fig.2A–B). We tested this hypothesis with a General Linear Model Hidden Markov Model (GLM-HMM), a newly established technique for identifying temporal patterns in behaviour^26,27^. GLM-HMMs extend the classical GLM by allowing parameter values to change over time-varying latent states. Transitions between latent states are governed by Markovian dynamics and are thus called ‘HMM-states’. In simple terms, thus, the technique amounts to decomposing behaviour with a series of GLMs with the same set of predictors, but in which the weights on predictors take different values at different points in time.

**Figure 2.**
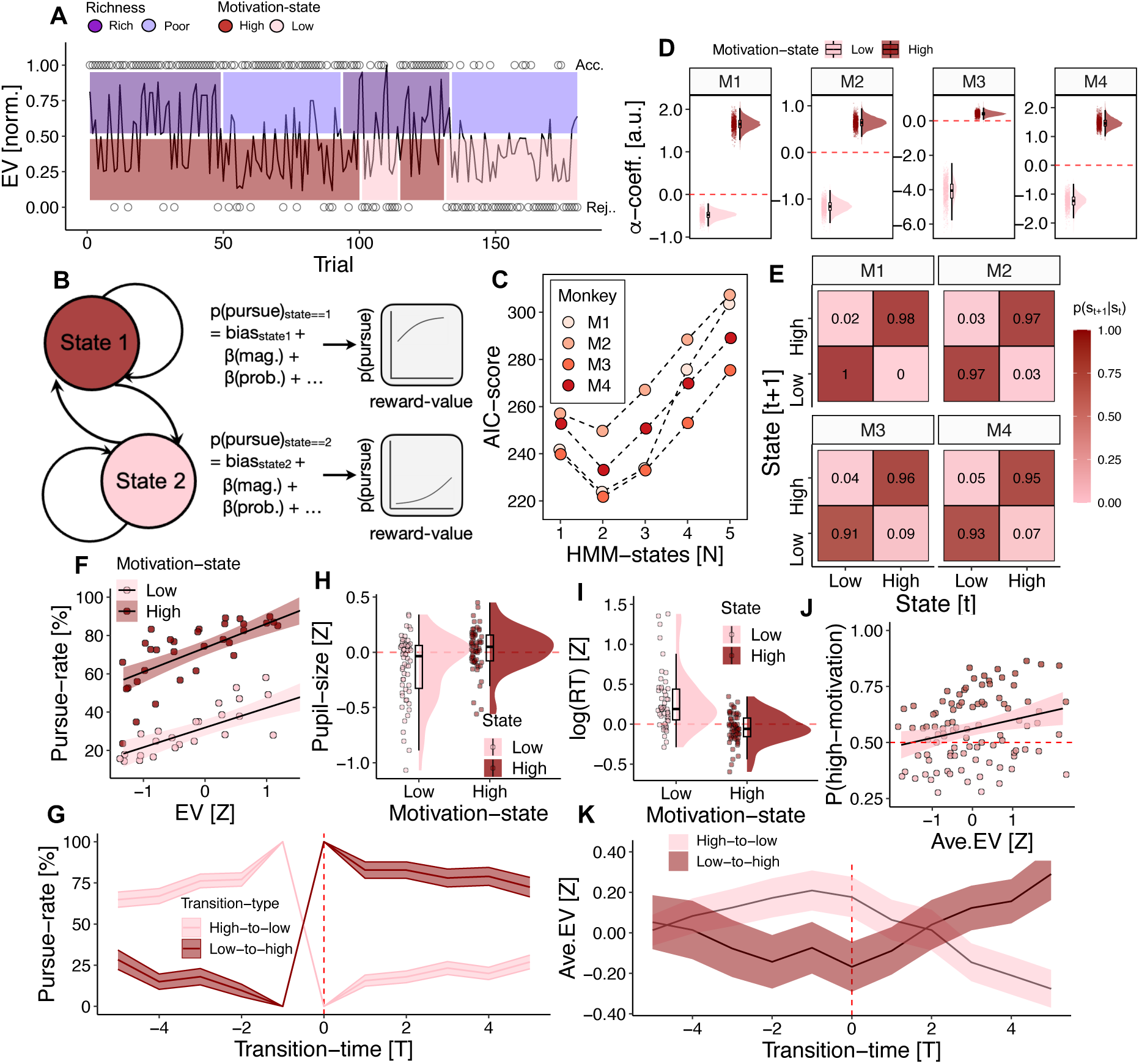
Animals exhibit distinct, persistent, and biologically meaningful motivation-states during behaviour. **(A)** Data showing reward-pursuit (dots linked to y-axis labels on right: dots at top indicate reward-pursuits; dots at bottom indicate reward-rejections) as a function of expectation-value (y-axis; left) over time (x-axis) in an example session. Animals exhibited autocorrelated patterns of behaviour in which they were biased to pursue (red; high-motivation) or reject (pink; low-motivation) opportunities almost regardless of expectation-value. States of high- and low-motivation (see panel B) tended to occur with experimentally controlled changes between rich (purple) and poor (blue) environments. **(B)** We captured such behaviour with a GLM-HMM featuring state-dependent bias parameters that produce corresponding state-dependent decision functions. These decision-functions capture changes in the way animals make decisions over time. **(C)** A model incorporating two HMM-states clearly improved model fit relative to alternative numbers of states. Reported here as AIC – a penalised form of log-likelihood (see supplementary fig.S2A for full cross-validation results). Points show mean session-wise AIC for each animal. **(D)** The posterior probabilities of state-specific bias parameters in all animals (M1 – M4) indicate that HMM-states impact an animal’s intrinsic propensity to pursue rewards. **(E)** Transition matrices reflecting the conditional probability of HMM-states from one trial to the next show strong autocorrelations in HMM-states in all animals (M1–M4). **(F)** Animals were more likely to pursue reward opportunities when occupying high-motivation states relative to low-motivation states regardless of an opportunity’s expected value. Dots indicate mean pursue-rates in quintiles of expected-value. **(G)** Pursue-rates changed abruptly at motivation-state transitions and are lower in the five trials before vs after low-to-high transitions and – correspondingly – greater in the five trials before vs after high-to-low transitions. This suggests that the GLM-HMM identifies state-like rather than gradual changes in behaviour. Color shades indicate transition direction. **(H)** Pupil-size – a common indicator of physiological arousal – is larger during the decision-phase (see Methods) of the task when animals are in the high-motivation state relative to the low-motivation state. Dots indicate mean pupil-size for each session in each animal in each motivation-state. In boxcharts, box-lengths indicate interquartile (IQR) ranges, bold lines indicate medians, and whiskers indicate median ± 1.5xIQR. **(I)** Reaction time – a common measure of vigour – is faster when animals occupy the high-motivation state. Dots indicate mean RT for each session in each animal in each motivation-state. In boxcharts, box-lengths indicate interquartile (IQR) ranges, bold lines indicate medians, and whiskers indicate median ± 1.5xIQR. **(J)** The likelihood that animals occupy the high-motivation state increases as a function of the expectation-value of recent reward opportunities (Ave. EV). Dots indicate mean rate of motivation-state occupancy in quintiles bins of average expectation-value. **(K)** Low-to-high and high-to-low transitions in motivation-states are consonant with changes in the availability of rewards. Lines and shadings indicate the mean and standard error, respectively, of Ave. EV in trials proximate to motivation-state transitions.

We formulated a GLM-HMM where decisions were predicted with binomial GLMs parameterised by: (i) a bias or ‘intercept’ term, (ii) a predictor for the expectation-value of the reward opportunity available on each trial, and (iii) predictors for external environment cues that were presented throughout the task (fig.2B; see methods for details). Per above, our focus was motivation-states corresponding to changes in an animal’s intrinsic drive for reward, as distinct from changes driven by external circumstances like the reward value at stake on a given trial. We therefore held the weights for predictors that described experimentally controlled aspects of the task constant and let only the bias parameters change between HMM-states. This configuration, we reasoned, was the best way of capturing autocorrelated patterns in behaviour whereby opportunities were pursued more frequently during periods of recent reward pursuit and accumulation.

We used a previously reported model-selection procedure which involved testing models with different numbers of HMM states using 5-fold cross validation^26^ (see supplementary figure S2A for details). We first validated the procedure using a series of parameter recovery tests (see supplementary figure S3A–C) before implementing it on behavioural data. This indicated that cross-validated log-likelihoods for 2-state GLM-HMMs were higher than 1-state GLM-HMMs for all animals (fig.2C; see supplementary figure S2A for full cross validation and S2B-D for further information), and we therefore used 2-state GLM-HMMs for all further analyses. Importantly, the 2-state GLM-HMMs for all animals featured clearly distinct state-specific bias parameters (fig.2D) and profoundly autocorrelated HMM-state transition matrices, which were consistent with state-like fluctuations in motivation for reward pursuit (fig.2E). Simulating data from fitted 2-state GLM-HMMs, moreover, reproduced key features of animal behaviour including richness of the environment and behavioural history effects (supplementary figure S2H–I), and qualitative patterns of autocorrelation in behaviour (supplementary figure S2E–G). Further analyses demonstrated that HMM-states were not simply reducible to satiety, fatigue, or time-on-task (S2B & S2D).

We tested the link between statistical HMM-states and biological motivation-states with a series of follow-up analyses. Firstly, we quantified differences in behaviour between HMM-states by decoding the *maximum a posteriori* HMM-state on each trial using a procedure that was validated on simulated data (correct HMM-state identified on 90% of trials; HMM-state transitions identified within +/− 3 trials of a true transition in 85% of cases; see supplementary figure S3). As expected, animals were markedly more likely to pursue reward opportunities in the putative high-motivation state compared to the low-motivation state regardless of the reward value available (GLM2.1; β_motivation-level_=2.64, *SE*=0.55, *p*<.001; fig.2F). Behaviour during transitions between HMM-states was characterised by abrupt changes in pursue-rates, suggesting that the GLM-HMM captured state-like differences in behaviour with a high degree of temporal precision (GLM2.2; β_before-vs-after-transition(high-to-low)_=-1.47, SE=0.09, *p*<.001; β_before-vs-after-transition(low-to-high)_=1.80, *SE*=0.30, *p*<.001, fig.2G). Finally, we established the model’s convergent validity by comparing HMM-states to: (i) trial-by-trial pupil-size, a well-validated indicator of physiological arousal^28^, and (ii) trial-by-trial reaction times (RT) on pursuit trials, a simple way of quantifying vigour. Consistent with the motivation-state view, putative high-motivation states featured faster RTs for pursue-decisions (GLM2.3; β_motivation-level_=-0.20, SE=0.03, *p*<.001; fig.2I) and increases in pupil-diameter during decision-making (GLM2.4; β_motivation-level_=0.20, SE=0.02, *p*<.001; fig.2H). The GLM-HMM, thus, provided quantitative evidence for discrete, persistent, and biologically meaningful internal motivation-states in monkey behaviour.

We next asked how motivation-states were related to the external environment: was it the case, for example, that an animal’s intrinsic motivation was shaped by the distribution of rewards in its milieu? We quantified the availability of rewards by calculating, for each trial, the average expectation-value of reward opportunities in the preceding five trials. Importantly, the expectation value of reward opportunities was experimentally controlled via the reward distributions characterising the task (fig.1B) and was therefore not affected by an animal’s decisions. Animals were, indeed, more likely to occupy high-motivation states as the expected-value of recent opportunities increased (GLM2.5; β_Ave.EV_=0.19, SE=0.09, *p*=.026; fig.2J) and visualising the expectation-value of rewards during transition points showed corresponding patterns, whereby high-to-low transitions occurred when average values decreased and vice versa for low-to-high transitions (fig.2K). The GLM-HMM therefore demonstrated not just that animals experienced state-like changes in motivation for rewards, but that these states were reconciled with the distribution of rewards in the world around them.

### Brain activity in DRN reflects an animal’s reward environment and changes in its motivation state

Animals performed the task under fMRI. Our analysis of fMRI recordings focused on *a priori* regions of interest (ROI; see fig. S4) comprising the ascending neuromodulatory systems (ANS) – an assemblage of phylogenetically ancient nuclei which includes the serotonergic DRN, in addition to the dopaminergic ventral tegmental area (VTA) and substantia nigra (SN), the cholinergic nucleus basalis (NB), and the noradrenergic locus coeruleus (LC). We also examined habenula (Hb) – an epithalamic nucleus with diverse subcortical connections that interact with the ANS in a reciprocal fashion^29,30^. Broadening our analysis beyond DRN provided a comparative perspective on its function, which is valuable given that reward functions are sometimes jointly attributed to different subcortical nuclei – the richness of the environment, for example, has been linked to both VTA and DRN related signals^19–21,31–37^, while various forms of reward uncertainty have been ascribed to DRN, LC and the cholinergic basal forebrain alike^15,38–42^. fMRI’s wholistic perspective enabled us to address this by comparing signals in DRN and other ANS nuclei^24^. This was accomplished with a novel suite of fMRI acquisition and pre-processing methods that optimised blood oxygen level dependent (BOLD) signal from subcortical regions and minimised artefacts and noise sources for midbrain and brainstem regions (see *Acquisition, reconstruction and pre-processing of MRI data* in Methods).

We first examined the links between brain activity and the reward environment during the pursue/reject decision made on each trial (GLM3.2; Methods). Due to the correlation between the richness of the environment and behavioural history, we separately analysed trials in which the previous encounter was pursued and rejected, respectively. This indicated that brain activity in DRN negatively correlated with the richness of the environment after rejection of opportunities (one-sample t-test; GLM3.2; *t_DRN; rejected_*(58)=-2.80, *p* =.034; *t_DRN; pursued_*(58)=-0.37, *p*=.713 after Bonferroni correction for multiple comparisons, as are all subsequent t-tests; fig.3A & 3B; see also fig.3A top panel for effect of richness of the environment in all trials; see fig.3 legend for further timing of BOLD signal). In contrast, VTA activity was prominent following opportunity pursuits while Hb exhibited aspects of both DRN and VTA patterns. Despite previous evidence linking DRN with reward uncertainty, we found no relationship between the environment’s stochasticity and brain activity in DRN (GLM3.1; *t_DRN_*(58)=1.10, *p*=.824) and no evidence that stochasticity was represented in other ROIs (GLM3.1; see fig.S5).

**Figure 3.**
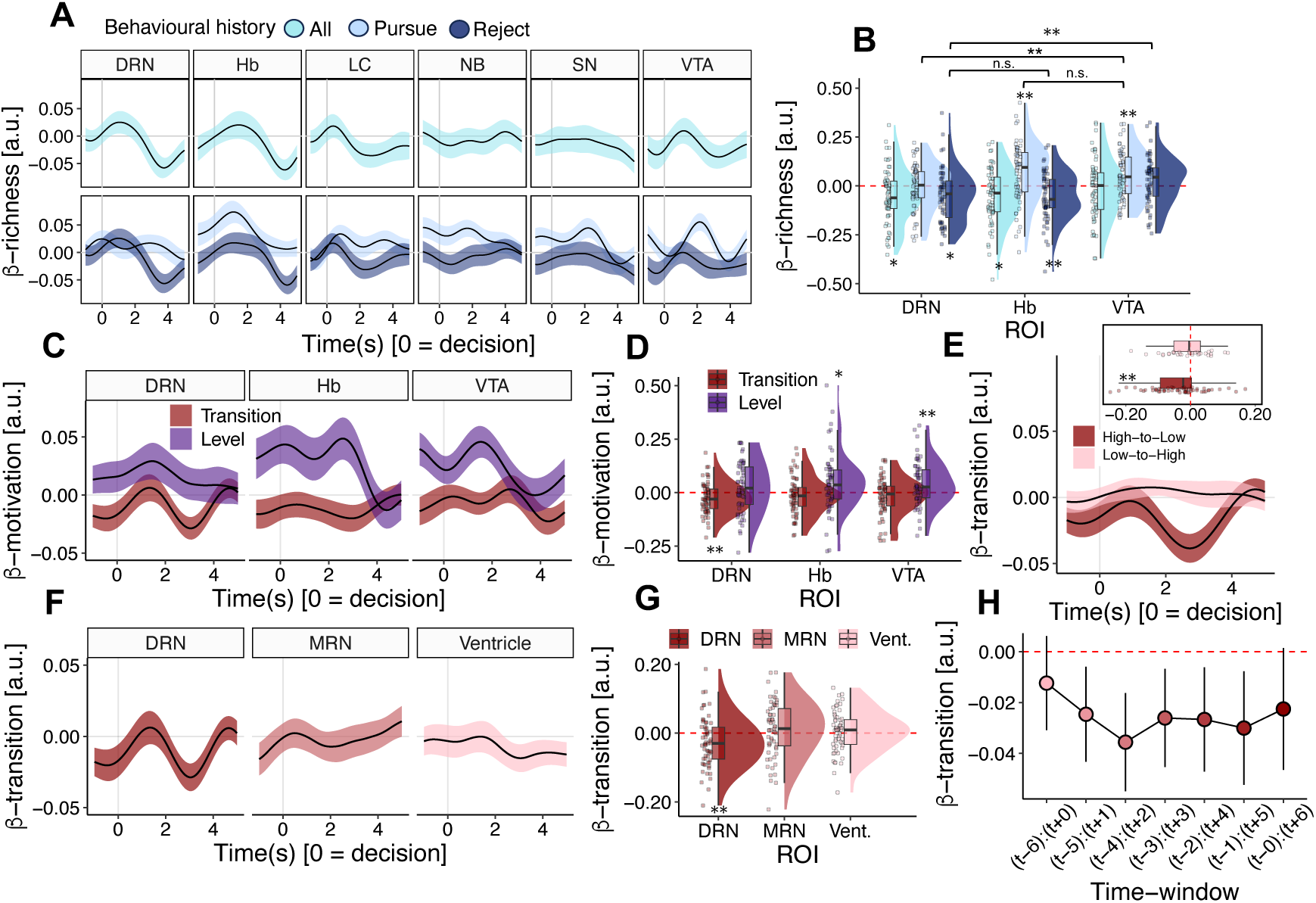
Brain activity in dorsal raphe nucleus represents a unique combination of an animal’s environment, recent behaviour and internal motivation-state. **(A)** Timecourse of the effect that the richness of the environment (β-richness) has on BOLD signal in subcortical ROIs. Time (x-axis) is depicted relative to the onset of reward-opportunities when pursue-vs-reject decisions are made (i.e. time=0 corresponds to decision-time). In macaque monkeys, BOLD signal reaches its maximum approximately 2-4s after the neural activity which caused it. The timing of peak β-weights in DRN, VTA and Hb is therefore consistent with brain activity that occurs at the time of decision-making. Top panel shows effects over all trials (GLM3.1) and bottom panel shows the analysis separately as a function of behavioural history (GLM3.2; see main text for explanation). Despite the partial collinearity between richness of the environment and behavioural history regressors, BOLD signal in both DRN and Hb represents richness of the environment across all trials (GLM 3.1; tDRN(58)=-2.79, p=.036; tHb(58)=-2.86, p=.035). **(B)** Distribution of peak effect sizes of richness of the environment (β-richness) on BOLD signal. Dots indicate peak effect sizes for individual sessions. Separating the data in this way revealed striking patterns in the direction and timing of environment representations across regions. DRN represented the richness of the environment with negative sign after the previous opportunity was rejected (GLM 3.2; tDRN; rejected(58)=-2.80, p=.034; tDRN; pursued(58)=-0.37, p=.713); BOLD signal in VTA represented richness with positive sign when animals pursued the previous opportunity – a signal that was hidden in the amalgamated data (GLM3.2; tVTA; pursued(58)=3.30, p=.009; tVTA; rejected(58)=1.60, p=.350); Hb, meanwhile, recapitulated the sign and behavioural contingency of DRN and VTA with distinct positive and negative signals on trials after pursuits and rejections, respectively (GLM3.2; tHb; pursued(58)=3.32, p=.009; tHb; rejected(58)=-3.37, p=.008; tHb; rejected-vs-pursued(58)=4.95, p<.001). A two-way ANOVA followed by pairwise comparisons confirmed that signals in DRN and VTA after rejections and pursuits, respectively, were different from one another but not from the corresponding Hb effects (FROI(2, 348)=8.106, p<.001, FBehav.-history(1, 348)=22.305, p<.001; FROI-by-Behav.-history(2, 348)=4.521, p=.012; tVTA vs DRN; pursued(58)=2.501, p=.015; tDRN vs VTA; rejected(58) =-3.403, p<.001; tVTA vs Hb; pursued(58) =0.650, p=.518; tDRN vs Hb; rejected(58)=-0.214, p=.831). **(C)** Timecourse of the effect of motivation-state level (high vs. low motivation state; purple) and motivation-state transitions (red) on BOLD activity in ROIs that represented the reward environment. Given haemodynamic delay (see **(A)**), the neural activity underlying the state-transition effect in DRN is likely to occur in the ITIs preceding trials on which transitions occur, meaning that it is not an epiphenomenon of action or outcome related processes. **(D)** Distribution of peak effect sizes for motivation-state level (purple) and motivation-state transitions (red) on brain activity. Only DRN represents transitions between motivation-states. Dots indicate peak effect sizes for individual sessions. **(E)** The timecourse of the effect that low-to-high (red) and high-to-low (pink) motivation-state transitions have on DRN BOLD activity. The effect is strongest for high-to-low state transitions. Inset panel shows distribution of peak effect sizes. **(F)** and **(G)** Motivation-state transitions affected BOLD signal in DRN but not neighbouring features like median raphe nucleus (MRN) and the 4^th^ ventricle, confirming that the pattern was DRN-specific (tDRN(58)=-3.69, *p*=.002, tMRN(58)=1.88, *p*=.102, tVent.(58)=-1.99, *p*=.102). **(H)** Shifting the position of transition-periods relative to decoded transitions showed that activity changed were temporally specific to decoded transition-times. X-axis indicates position of transition-period relative to decoded transition time (where t=0 represents decoded transition trial). Dots and error bars indicate mean and 95% confidence interval of effect-sizes for each transition-window. See supplementary figure S8 for further detail. In timecourse graphs (A, C, E and F), lines and shadings show the mean and standard error (SE) of the β weights across the sessions, respectively. Distribution graphs (B, D, E inset, and G) show peak regression weights where dots indicate weights from individual sessions. Dots and errorbars in H show mean and 95% confidence interval of regression effects for different transition-period windows. In boxcharts, box-lengths indicate interquartile (IQR) ranges, bold lines indicate medians, and whiskers indicate median ± 1.5xIQR.

After determining that DRN, VTA and Hb were the key regions tracking the environment, we probed their involvement in motivation-states identified with the GLM-HMM. We first quantified the relationship between brain activity and motivation-state *level* (high vs. low) which suggested that high motivation-states were reflected in increased activity in VTA and Hb, but not in DRN (GLM3.4; *t_Hb; motivation-state_* (58)=2.41, *p*=.038; *t_VTA; motivation-state_* (58)=2.96, *p*=.014; *t_DRN; motivation-state_* (58)=1.59, *p*=.120; fig.3C & fig.3D). Next, we explored activity related to motivation-state *transitions* (change vs. no-change) by identifying transition-trials on which the *maximum a posteriori* motivation-state was different from the preceding trial. For each transition-trial, we defined a corresponding transition-period comprising a symmetric 7 trial window centred on the transition-trial (i.e. the transition trial +/− 3 trials) – a timespan that was based on simulations showing that transition decoding was accurate at this timescale (see supplementary figure S3F). Brain activity in these periods showed a striking dissociation whereby DRN – but neither Hb nor VTA – signalled transitions between motivation-states (GLM3.4; *t_DRN; state-transition_*(58)=-2.70, *p*=.020; *t_Hb; state-transition_*(58)=-1.28, *p*=.206; *t_VTA; state-transition_*(58)=-1.65, *p* =.206; fig.3C & 3D).

We confirmed that the transition effect was localised to DRN by analysing BOLD signal from adjacent anatomical features such as the medial raphe nucleus (MRN) and fourth ventricle, which showed no corresponding patterns (fig.3F & 3G; t_DRN_(58) =-2.70, *p*=.020, t_MRN_(58)=0.93, *p*=.715, t_Vent._(58)=0.54, *p* =.715). We then systematically varied the position of transition-periods relative to transition-trials to probe the effect’s temporal specificity. Activity changes in DRN were most prominent when the analysed period was closely aligned with the decoded transition-trial, suggesting that they were indeed specific to motivation-state transitions (fig.3H; see Supplementary fig.S8B & S8C for details). Given the haemodynamic delay in fMRI recordings, the timing of the effects suggested that activity changes occurred during the ITIs preceding trials on which state-transitions occurred, meaning that they are unlikely to be epiphenomena of action or outcome-processing events (supplementary fig.S8B). Finally, we distinguished different directions of transition which indicated that DRN activity covaried with high-to-low transitions but not low-to-high transitions (GLM3.4 *t_DRN; high-to-low-transitions_* (58)=-2.46, *p*=.017; *t_DRN; low-to-high-transitions_* (58)=1.24, *p*=.220; fig.3E). Taken together, these analyses suggested that DRN – and DRN specifically – implemented negative changes in intrinsic motivation for rewards.

### Non-invasive disruption of DRN perturbs motivation-state transitions

fMRI recordings linked DRN to behaviour in two complementary ways; (i) DRN coded the richness of the environment, which modulated an animal’s motivation to pursue specific reward opportunities and (ii) DRN coded transitions between GLM-HMM derived motivation-states, and therefore changes in reward pursuit over multi-trial timescales. We tested DRN’s causal contribution in these respects with a second experiment using transcranial ultrasound stimulation (TUS; fig.4A) – a minimally invasive and reversible technique that disrupts brain activity via kinetic interactions between focused ultrasound waves and mechanosensitive ion channels at the neuron and astrocyte membrane^43,44^. These interactions change activity in spatially circumscribed brain regions, and short TUS trains of the type we used are known to produce short term changes in neural activity by inducing N-methyl-d-aspartate (NMDA)-dependent plasticity^45–49^. As a result, a region targeted with TUS alters its responsiveness to activity in interconnected areas, while non-stimulated areas show no such change^25^.

**Figure 4.**
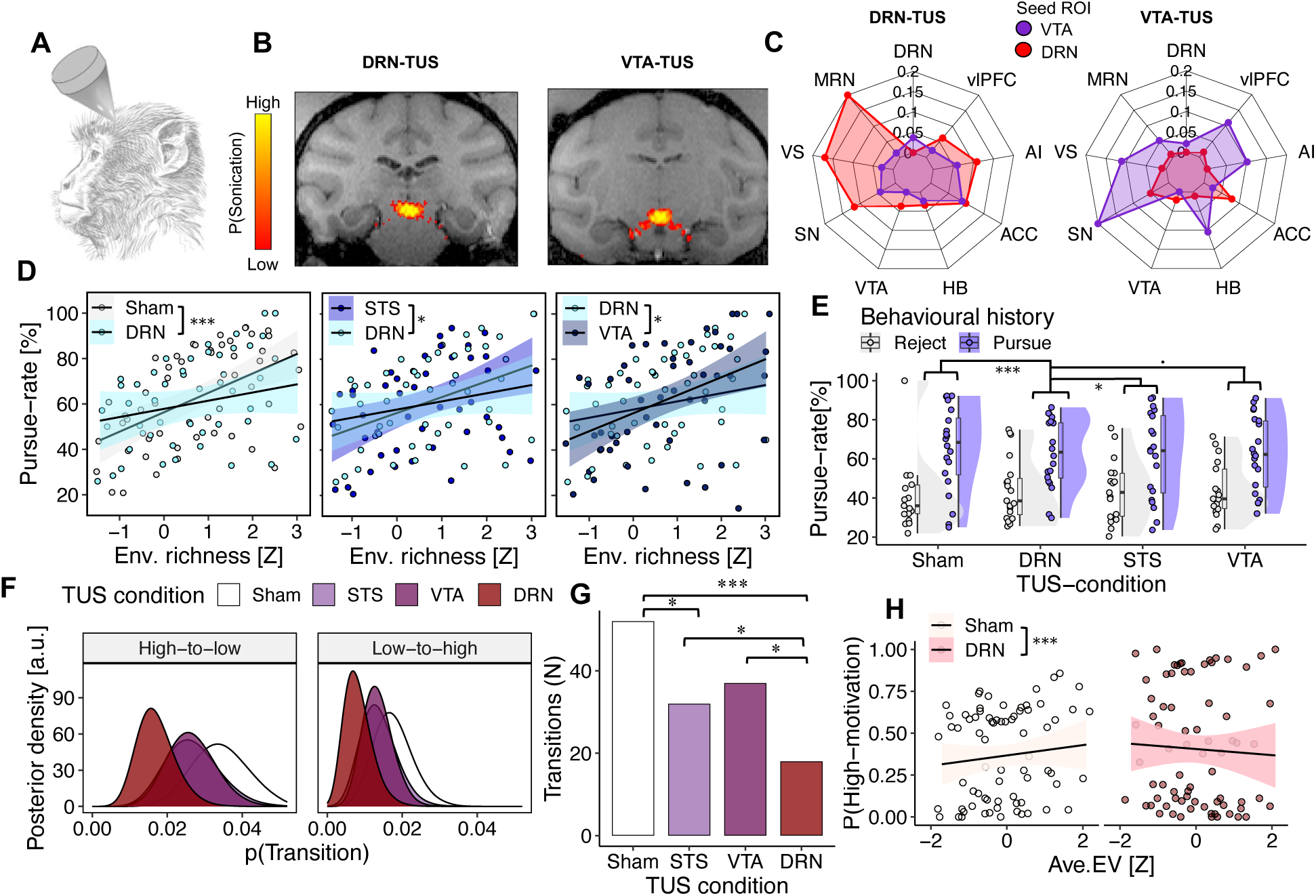
Non-invasive perturbation of dorsal raphe nucleus shows that it is causally involved in the influence of a monkey’s environment on its behaviour. **(A)** We investigated the causal contribution of DRN to behaviour with transcranial ultrasound stimulation (TUS) – a minimally invasive form of brain stimulation suited to subcortical brain regions **(B)** Simulating the propagation of acoustic waves under the TUS protocol suggested that sonication was target-specific after DRN-TUS and VTA-TUS. Colour map indicates probability of neuromodulation after bilateral TUS, calculated as the sum of stimulation intensity (Isppa) maps across two consecutive stimulations delivered over left and right hemispheres, respectively (see fig. S9). Combined impact probability maps are overlaid on a standard F99 brain. **(C)** DRN-TUS (left) disrupted DRN (red) connectivity with key brain regions whilst leaving VTA (blue) connectivity intact. The converse pattern occurred after VTA-TUS (right). Radial axis shows absolute-value of the difference in connectivity between seed and target ROIs pre-vs-post TUS (see methods for details). Targets included ventrolateral prefrontal cortex (vlPFC), anterior insular (AI), anterior cingulate cortex (ACC), habenula (Hb), ventral tegmental area (VTA), substantia nigra (SN), ventral striatum (VS), median raphe nucleus (MRN) and dorsal raphe nucleus (DRN). **(D)** DRN-TUS (left) attenuated the effect that the richness of an animal’s environment had on behaviour relative to sham-TUS (left), STS-TUS (middle), and VTA-TUS (right). Datapoints indicate mean pursue-rates in quintile bins of richness-of-the-environment. **(E)** DRN-TUS attenuated the effect of behavioural history on current behaviour relative to sham-TUS and STS-TUS, marginally with respect to VTA-TUS. Datapoints indicate mean pursue-rates for each animal in each session for. In boxcharts, box-lengths indicate interquartile (IQR) ranges, bold lines indicate medians, and whiskers indicate median ± 1.5xIQR. **(F)** The posterior-probability of state-transitions in GLM-HMMs fitted to data from different TUS conditions. The central tendency of DRN-TUS posteriors is lower than for to STS-TUS, VTA-TUS and sham-TUS. (**G)** Fewer motivation-state transitions occur in DRN-TUS sessions relative to all other TUS conditions. Bars show the total number of motivation-state transitions decoded over all sessions in each TUS condition. **(H)** DRN-TUS diminishes the effect that the availability of rewards ordinarily has on an animal’s motivation-state level. Datapoints indicate mean rates of high-motivation state occupancy in quintiles of average expected value (Ave. EV).

**Figure 5.**
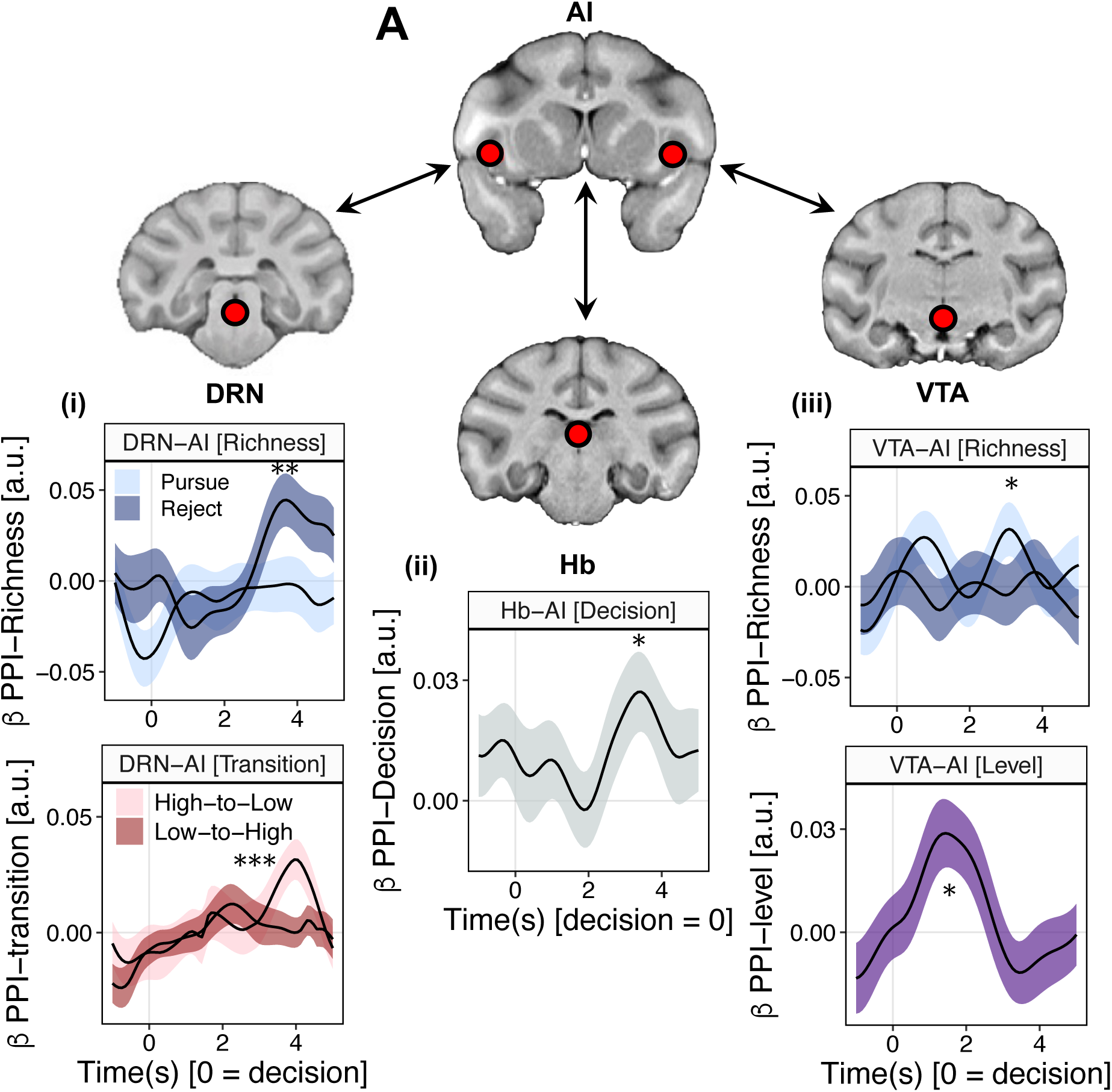
A brain circuit for reconciling an animal’s motivation with its environment. **(A)** PPIs indicated the following patterns of cortico-subcortical. **(i**) Connectivity between DRN and AI increased as a function of richness of the environment in the aftermath of rejections (GLM3.6; upper panel), and as a function of high-to-low motivation-state transitions (lower panel; GLM3.7; tPPI(high-to-low)=3.32, *p*=.002, tPPI(low-to-high)=1.10, *p*=.274), consistent with DRN’s fMRI signals **(ii)** connectivity between AI and Hb increased as a function of an animal’s ultimate decision about whether to pursue a reward opportunity (GLM3.8). **(iii)** In contrast to DRN, connectivity between VTA and AI increased as a function of richness of the environment in the aftermath of pursuits (GLM3.6; upper panel), and during high-motivation states (GLM3.7; tmotivation-level=2.26, *p*=.028).

We used an offline TUS protocol that modulates neural activity for over an hour after sonication^25,50,51^. We compared DRN-TUS to three control conditions: a sham condition (sham-TUS), an active cortical control (superior temporal sulcus; STS-TUS), and an additional active subcortical control in VTA – a region implicated in motivation in fMRI recordings but in different ways to DRN. We first performed simulations of acoustic wave propagation under the TUS protocol which established that selective perturbation of DRN and VTA was feasible despite their diminutive size and anatomical location (fig.4A & 4B, and supplementary figure S9). We then empirically tested the TUS protocol’s anatomical specificity by measuring changes in functional connectivity between DRN, VTA and series of key brain areas before-vs-after DRN-TUS and – as a control – before-vs-after VTA TUS. This showed striking and selective changes in region-specific functional connectivity: DRN-TUS disrupted DRN’s ordinary patterns of coactivation with key interconnected regions but left VTA connectivity unchanged, and the converse pattern occurred after VTA-TUS (fig.4C). In combination with the sonication simulations, this indicated that TUS induced, safe, efficacious, and highly localised disruption of DRN and VTA activity, analogous to previous investigations of TUS in subcortical brain regions^25,50,51^.

We tested the same monkeys from the fMRI experiment on five sessions of the behavioural task per TUS condition in counterbalanced and pseudorandomised order (see Methods). We first investigated how TUS modulated the influence of (i) richness of the environment, and (ii) behavioural history, which were the key factors driving the pursue/reject decisions made on each trial. DRN-TUS diminished the effect of richness of the environment relative to all other conditions. (GLM4.1; β_DRN-TUS vs sham-TUS*richness_=-0.18, SE=0.05, *p*<.001; β_DRN-TUS vs STS-TUS*richness_=-0.12, SE=0.06, *p*=.025; β_DRN-TUS vs VTA-TUS*richness_=-0.12, SE=0.06, *p*=.035; fig.4D). Similarly, DRN-TUS clearly reduced the effect of behavioural history relative to sham-TUS and STS-TUS, and produced a marginal difference relative to VTA-TUS (GLM4.2; β_DRN-TUS vs sham-TUS*behave-history_=-0.42, SE=0.11, *p*<.001; β_DRN-TUS vs STS-TUS*behave-history_=-0.24, SE=0.11, *p*=.027; β_DRN-TUS vs VTA-TUS*behave-history_=-0.20, SE=0.11, *p*=.057; fig.4E). There were no differences between active control conditions (STS-TUS and VTA-TUS) and sham-TUS (GLM4.1 & fig.4D; β_VTA-TUS vs sham-TUS*richness_=-0.06, SE=0.06, *p*=.282; β_STS-TUS vs sham-TUS*richness_=-0.06, SE =0.05, *p*=.306; GLM4.2 & fig.4E; β_VTA-TUS vs sham-TUS*behavioural-history_=-0.20, SE=0.10, *p*=.060; β_STS-TUS vs sham-TUS*behavioural-history_=-0.18, SE=0.11, *p*=.090). DRN, therefore, was specifically and causally involved in the impact of an animal’s recent reward environment and behaviour on its present decisions to pursue reward.

Next, we applied the GLM-HMM to behavioural data from the TUS experiment (see Methods). Our analysis focused on motivational-state transitions, which were a distinguishing feature of DRN activity in fMRI recordings. These recordings suggested that DRN was specifically involved in high-to-low motivational-state transitions, but we first performed the analysis in a direction-neutral way given the interdependence of time-series observations: because animals typically began the task in a high-motivation state (p(init-state=high)=0.70), reducing the likelihood of high-to-low transitions would necessarily reduce the frequency of low-to-high transitions in a fixed-length time series, even if the underlying low-to-high transition probability was unchanged. The analysis showed a striking pattern whereby DRN-TUS reduced the likelihood of transitions relative to all other TUS conditions (GLM 4.3; β_DRN-TUS vs sham-TUS_=-1.08, SE=0.27, *p*<.001; β_DRN-TUS vs VTA-TUS_=-0.71, SE=0.29, *p*=.013; β_DRN-TUS vs STS-TUS_=-0.57, SE=0.29, *p*=.051; fig.4F). There was no effect of VTA-TUS relative to sham-TUS (GLM4.3; β_VTA-TUS vs sham-TUS_ = −0.37, SE = 0.22, *p* = .090). There was a moderate difference between STS-TUS and sham-TUS but this was outweighed by the fact that DRN-TUS reduced the likelihood of transitions even relative to STS-TUS (GLM4.3; β_STS-TUS vs sham-TUS_=-0.51, SE=0.23, *p*=.022; β_DRN-TUS vs STS-TUS_=-0.57, SE=0.29, *p*=.051). DRN-TUS was, moreover, the only condition that reduced high-to-low transitions specifically (GLM4.4; β_DRN-TUS vs sham-TUS_=-0.77, SE=0.32, *p*=.016; β_VTA-TUS vs sham-TUS_=-0.33, SE=0.28, *p*=.238; β_STS-TUS vs sham-TUS_=-0.43, SE=0.29, *p*=.139), and caused animals to spend more time in high-motivation states relative to sham-TUS (GLM4.5; β_DRN-TUS vs sham-TUS_=0.28, SE=0.07, *p*<.001), consistent with the idea that DRN-TUS prevented animals from transitioning to low motivation states.

Finally, we sought to specify how DRN controlled transitions between motivation states. To do so, we returned to an earlier analysis demonstrating that motivation-states were consonant with the availability of rewards: in brief, animals were more likely to occupy high-motivation states when there were many high-value rewards available, analogous to the way that animals pursued more opportunities in rich environments (fig.2J & 2K). Given DRN’s critical involvement in the latter phenomenon, we asked whether its control of motivational-state transitions was mediated by the availability of rewards. Consistent with this view, DRN-TUS diminished the relationship between the availability of rewards and motivation-states relative to all other TUS conditions (GLM4.5; fig.4H; β_DRN-TUS vs sham-TUS*Ave. EV_=-0.28, SE=0.07, *p*<.001; β_DRN-TUS vs STS-TUS*Ave. EV_=-0.29, SE=0.07, *p*<.001; β_DRN-TUS vs VTA-TUS*Ave. EV_=-0.30, SE=0.07, *p*<.001). There were no effects of VTA-TUS or STS-TUS relative to sham-TUS (β_VTA-TUS vs sham-TUS*Ave. EV_=-0.03, SE=0.06, *p*=.610; β_STS-TUS vs sham-TUS*Ave. EV_=-0.05, SE=0.07, *p*=.499). Intriguingly, DRN-TUS’s influence was more prominent for the link between low-value reward environments and low-motivation states, which dovetailed with fMRI signals implicating DRN specifically in high-to-low transitions (GLM4.5; for Ave.EV< μ_Ave.EV_ β_DRN-TUS*Ave. EV_=-0.52, SE=0.19, *p*=.006; for Ave.EV ≥ μ_Ave.EV_ β_DRN-TUS*Ave. EV_=-0.13, SE=0.15, *p*=.403). Taken together, these results suggest that DRN has a fundamental role in ensuring that an animal’s motivation-level is appropriate to the distribution of rewards in the environment.

### A cortico-subcortical circuit for reconciling behaviour with the environment

In a final analysis, we examined interactions between DRN and other brain regions that might relate to its behavioural function. To do so, we first reverted to the analysis of brain activity and behaviour which did not rely on the GLM-HMM framework (fig.3A & B). Here, DRN, VTA and Hb represented an animal’s environment with complementary patterns of activity, which suggested that they might form a circuit for reconciling decisions with the surrounding environment. In testing the circuit hypothesis, we expanded our purview to functionally related cortical ROIs in Supplementary Motor Area (SMA), Anterior Cingulate Cortex (ACC) and Anterior Insula (AI) based on previous work implicating these regions in behavioural change^19,50,52–54^. Only AI recapitulated the distinctive contingency between richness and behavioural history seen in subcortical ROIs, and we therefore retained AI as a cortical ROI for connectivity analysis (see fig.S7).

Next, we asked which ROIs coded the pursue/reject decision taken on each trial – in other words, the output of the putative decision-making circuit. BOLD activity time-locked to decision-making in both Hb and AI represented pursue/reject decisions (GLM3.3; *t_pursue; Hb_*(58)=3.24, *p*=.011; *t_pursue; AI_*(58)=2.78, *p*=.028; see fig.S6). We then conducted psychophysiological interaction (PPI) analyses to probe changes in pairwise connectivity between regions as a function of the environment^55,56^. PPIs mirrored the patterns observed earlier: connectivity between DRN and AI increased as a function of richness of the environment after *rejections* (GLM3.6; *t_PPI(DRN-AI by richness); rejected_*(58)=2.84, *p*=.006; fig.5Ai) and increased between VTA and AI after *pursuits* (GLM 3.6; *t_PPI(AI-VTA by richness); pursued_*(58)=2.14, *p*=.036; fig.5Aiii). Adopting the GLM-HMM analysis approach showed a corresponding pattern of results (GLM3.7; figs.5Ai & 5Aiii). Connectivity between Hb and AI was not modulated by the richness of the environment but instead by the pursue/reject decision taken on each trial, consistent with a role in translating motivation to action^57^ (GLM3.8 *t_PPI(AI-Hb by action)_*(58)=2.25, *p*=.028; fig.5Aii).

## Discussion

We provide converging evidence that DRN controls changes in an animal’s motivation for rewards. We observed distinctive patterns of DRN activity corresponding to the richness of an animal’s environment and transitions between statistically delineated motivation-states. We followed this with first-of-its-kind minimally invasive DRN disruption which diminished the environment’s effect on decision-making and reduced the frequency of motivation-state transitions. This suggests that DRN is causally involved in changes in motivation.

Our findings dovetail with two prevailing views on DRN function: the first has emphasised DRN’s role in behavioural changes, for example between exploration and exploitation, or patience and impulsivity^7–14^; the second has linked DRN activity to statistical descriptions of rewards, like the general value and/or uncertainty of the options characterising an environment^15–23^. We observed similar brain-behaviour relationships here, and the concurrence of both phenomena suggests that they are fundamentally related. In support of this view, we found that DRN disruption impaired animals from matching their motivation-state with the availability of rewards, and specifically from entering low-motivation states during low-value environments – an important motif in normal behaviour. In tandem with previous studies, this indicates that DRN is critical for changing an animal’s behaviour according to the reward statistics of the surrounding world.

Several previous studies have linked DRN with prediction-error like responses during learning tasks in which there is fundamental uncertainty about the reward available on each trial^7,9,11,15,38,58^. In contrast, the properties of specific reward opportunities were explicitly signalled in our paradigm, and the key manipulation was the distribution of reward opportunities over time. This might explain the absence of reward-uncertainty signals in DRN fMRI recordings. It might similarly explain VTA’s involvement in the task. Although it is perhaps surprising that VTA did not feature more prominently given its well-documented links to reward and motivation, the task did not require reinforcement learning, which is a key function of dopaminergic nuclei ^59^. The patterns of VTA activity that we did observe – (i) the richness of the environment after pursuits, and (ii) trial-by-trial motivation-state level – are consistent with a cognate role in driving behaviour in proximity to rewards^31,36,37,60^.

How might DRN coordinate with other brain regions to control decisions? Unlike other subcortical ROIs, we found that activity in Hb covaried with both an animal’s pursue/reject decisions and the individual factors that shaped them. This is consistent with an emerging theory that Hb integrates motivationally-salient information from its diverse range of afferent connections to control motor output via the basal ganglia^57,61–64^. Similar patterns of activity occurred in AI, which is perhaps the computational source of representations concerning rewards obtained over multiple timepoints^19,57,65^. Although it is difficult to specify directions of influence from fMRI recordings, we observed patterns of functional connectivity consistent with communication between AI-DRN, AI-VTA, and AI-Hb depending on an animal’s previous behaviours – a scheme which matches well-documented patterns of anatomical connection^62^. Cortico-subcortical interactions of this kind echo a common neural motif whereby information synthesised in the cortex is transmitted to subcortical nuclei, which have wide-ranging efferent connections capable of orchestrating brain-wide activity^57,63,66,67^. The fact that animals were *ceteris paribus* more likely to pursue rewards in high-value environments controverts prominent formulations of optimal reward-seeking^1,19,31,50,65^, which argue that animals should be less selective during periods of reward scarcity on account of their homeostatic requirements. Nevertheless, the pattern observed ensures that animals maximise their reward intake in the manner that these theories envisage: given that animals usually lack perfect knowledge about the richness or sparseness of future opportunities, a reasonable strategy is to concentrate their efforts on rich environments where they have the most to gain. In support of this view, the GLM-HMM analysis identified persistent motivation-states in monkey behaviour that were positively correlated with the availability of rewards.

Finally, it is noteworthy that many neurons in DRN are serotonergic and that DRN is the principal source of serotonin to the mammalian forebrain^3,4^. Although the techniques we implemented are not serotonin specific, our results are reminiscent of several proposals about its function. We found (i) a positive correlation between DRN activity and reward scarcity, evoking classic theories that link serotonin to punishments and reward omissions^32,68–70^, and (ii) a causal relationship between DRN and changes in behaviour, akin to serotonin’s role in perseverative choice^7,11,13,71^. The key finding in our experiment relative to previous work is that DRN controlled patterns of behaviour that unfolded over multiple timepoints, not just specific reinforcement events or decisions. In a similar vein, it is possible that serotonin is important not for moment-to-moment decision-making *per se* but for changing the overarching strategy of an animal’s behaviour, and perhaps especially for changes that involve adverse feedback from the environment. Testing this hypothesis is a promising avenue for future experiments.

## Data and code availability

Data and analysis scripts supporting the findings of this manuscript will be available online upon acceptance for publication. Additional information can be accessed upon request from corresponding author.

## Acknowledgements

The authors thank the Biomedical Services staff at the University of Oxford for their care of the macaques. The work was funded by the BBSRC (BB/W008947/1; BB/W003392/1), Wellcome Trust (221794/Z/20/Z; 203139/Z/16/Z). and Medical Research Council (MR/P024955/1).

## Author contributions

LP, NK and MSFR conceived the study. LP and NK performed the fMRI experiments. MS and MC developed the fMRI acquisition protocol and pre-processing pipeline. NK, XC and RC prepared and validated the TUS equipment. LP and NK performed the TUS experiments. LP, NK, AM and MSFR analysed the data. LP wrote the first draft of the manuscript. LP, NK and MSFR contributed to subsequent versions of the manuscript. All authors approved the final manuscript.

## Competing interests

the authors declare no competing interests.

## Additional information

## SUPPLEMENTARY MATERIAL

**Figure S1.**
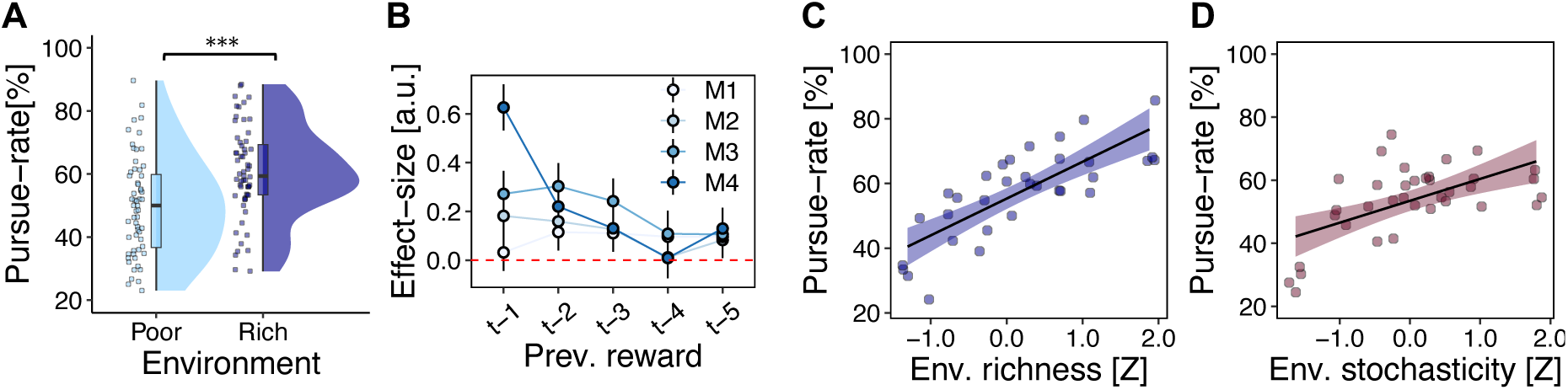
Additional analysis of behaviour. **(A)** A complementary perspective on the relationship between environments and behaviour is afforded by the blocks specified in the experimental design. The reward-probability distributions for rich-volatile and poor-volatile blocks partly overlapped (fig.1B), which produced a subset of trials with the same reward value in different contexts (poor vs rich). Animals were more likely to pursue offers that occurred in rich blocks relative to poor blocks, dovetailing with the analysis reported in the main text (fig.1E; β_rich-vs-poor_=0.90, *SE*=0.14, *p*<.001). Dots indicate mean rates of responding in each session and animal. In boxcharts, box-lengths indicate interquartile (IQR) ranges, bold lines indicate medians, and whiskers indicate median ± 1.5xIQR. **(B)** In the main text, we operationalised reward environments in way that emphasised an animal’s recent experience reward outcomes, rather than underlying experimentally defined conditions. We set the time-horizon for this analysis by testing the effect of specific reward outcomes received *n* trials into the past. Rewards received 5 trials into the past (*t*-5) were the last point at which significant effects occurred in all animals, and we therefore operationalised environment-related effects with 5-trial retrospective windows. Points and error bars indicate mean and 95% confidence interval for effect-sizes of previous rewards for each individual animal. **(C) and (D)** Animals were more likely to pursue offers as a function of increases in the richness **(C)** and stochasticity **(D)** of the environment (see also fig.1E–F). Dots indicate mean pursue-rates in quintile bins of richness-of-the-environment and stochasticity-of-the-environment, respectively.

**Figure S2.**
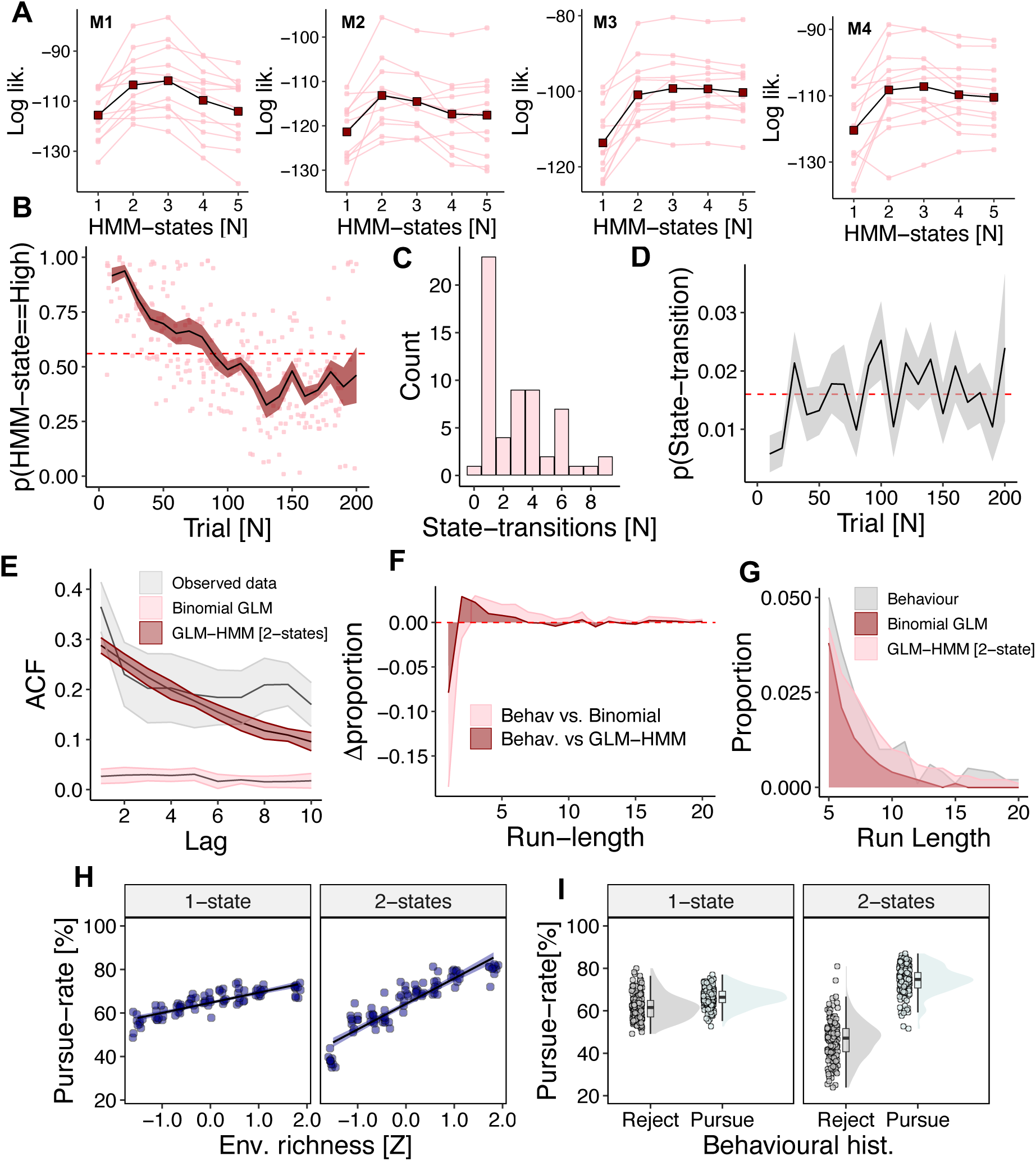
Quantitative and qualitative validation of the GLM-HMM. **(A)** We tested GLM-HMMs using 5-fold cross-validation performed within each individual animal. This involved dividing the data into five folds and iteratively fitting the model to sessions in four folds (i.e. training folds) before testing performance on the held-out (i.e. test) fold. Our metric of model performance was log-likelihood, which we computed using the forward algorithm (see Methods). We constructed folds between-sessions instead of trials-within-sessions because between-session variance is greater than within-session variance ^26^, meaning that between-session cross-validation is a more demanding tests of performance. For each fold, we tested models with *s* ∈ {1, 2, 3, 4, 5} HMM-states and calculated the log-likelihood of test sessions. All animals followed a similar pattern whereby models with two HMM-states performed better than Binomial GLMs (i.e. 1-state models), whereas additional states either failed to improve or impaired model performance. The relationship between the log-likelihood of the held-out sessions as a function of HMM-states is shown for all animals (foregrounded bold lines indicate mean log-likelihood of held-out sessions; background lines indicate log-likelihood of individual sessions). **(B)** The relative probability of motivation-states (y-axis) over time (x-axis). Animals are more likely to occupy high-motivation states early in a session, and approximately equally likely to occupy high-vs-low motivation states at the end of a session – motivation-state, thus, are not reducible to satiety, fatigue, or time-on-task. Black line and shaded areas indicate mean and SEM, respectively, of state-occupancy as a function of time; dots indicate mean levels of state-occupancy in quintile time-bins over individual sessions. Dashed red line indicates mean probability of motivation-state occupancy, regardless of time. **(C)** Histogram of the number of motivation-state transitions per session (x-axis). One session features no transitions. Data from all animals included. **(D)** The probability of motivation-state transition (y-axis) as a function of time (x-axis). Although transitions are more likely to occur in later trials, there is no consistent and specific temporal pattern in transition events indicating that they are not reducible to the passage of time (per (**C)**). Line and shaded area indicate mean and SEM, respectively, of state-transitions rates in quintile bins of trials. Dashed red line indicates overall mean probability of state-transition, regardless of time. Data from all animals included. **(E)** Simulated datasets from fitted 2-state GLM-HMMs reproduce trial-to-trial autocorrelations in behaviour, which were a key feature of animal behavioural. A binomial GLM – in contrast – is unable to reproduce this pattern. X-axis shows autocorrelation function (ACF) between the decision on trial *t* and decisions on (*t-1):(t-10)* for recorded behaviour, the 2-state GLM-HMM and a Binomial GLM for an example animal. Lines and shaded areas show the mean and SEM, respectively, of the ACF at each individual lag. ACFs for the behavioural data were obtained by calculating the ACF at each lag in each individual session (N=16) and taking the lag-wise mean and SE of ACFs. Simulated ACFs were obtained by fitting the GLM-HMM using the 5-fold cross validation method described above and simulating decisions on held-out sessions. 10 simulations of each held-out session were performed before calculating the lag-wise mean and SE of the resulting distribution of ACFs. **(F)** A transparent way of showing autocorrelations in behaviour is with decision run-lengths – that is, the number of times that the same decision is repeated over consecutive trials (rejecting three consecutive reward opportunities, for example, constitutes a decision run-length of 3; see also^26^). Comparing the relative proportion of run-lengths in simulated and behavioural data showed that the GLM-HMM captured animal behaviour better than a binomial GLM. Data shown as difference in observed and simulated (behaviour – model simulations) in proportion of run-lengths for the same example animal and simulations as in panel (E). **(G)** In particular, the GLM-HMM captures the tendency of animals to repeat decisions over five or more consecutive trials. A binomial GLM, in contrast, was unable to reproduce this pattern. Data from the same example animal and simulations as panel (E). **(H)** and **(I)** Fitting the GLM’s (GLM 1.1–1.4) used to analyse behaviour to data that was generated from a 2-state GLM-HMM recapitulated richness of the environment **(H)** and behavioural history effects **(I)**. The same patterns did not occur for data simulated from 1-state (i.e. binomial) GLMs. See fig. 1G and supplementary figure S1C for comparison.

**Figure S3.**
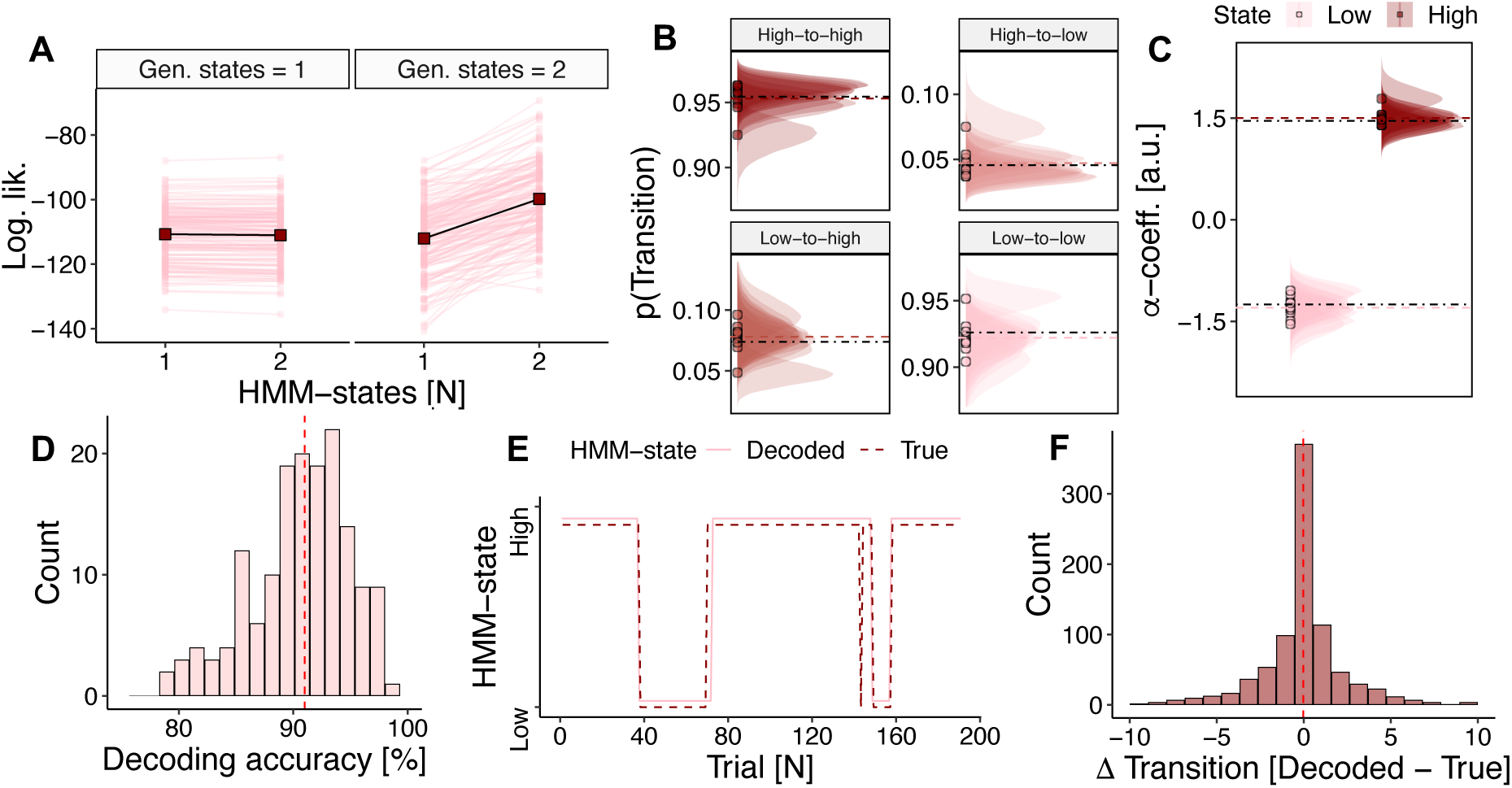
Parameter recovery and decoding accuracy of GLM-HMM. We assessed the performance of the GLM-HMM fitting procedure by simulating 10 datasets from fitted GLM-HMMs. Each dataset was approximately the same size as the total data obtained from each individual animal (∼16 sessions, ∼3000 trials). **(A)** We first tested whether the fitting procedure recovered the true number of HMM-states when the data was generated by a 1-state (i.e. Binomial GLM; left panel) and 2-state GLM-HMM (right panel) models. 2-state GLM-HMMs performed better under 2-state but not 1-state generative processes. All subsequent data pertains to simulations with 2-state GLM-HMM generative models. **(B)** The fitting procedure successfully recovered generative transition-matrix parameters. Graphs show posterior distributions over transition-matrix components in fitted 2-state GLM-HMMs. Black dashed lines indicate generative parameters and coloured lines indicate mean parameters over all fitted models. **(C)** The fitting procedure successfully recovered state-specific bias/intercept parameters. Graphs show posterior distributions over state-specific bias values. Dashed black lines indicate generative parameters and coloured lines indicate mean parameters over all fitted models. **(D)** We tested the accuracy of the Viterbi decoding algorithm by applying it to simulated datasets in which the true HMM-state was known. Viterbi decoding was successful on 90.68% (± 0.72%) of trials. Histogram shows session-level decoding accuracy (%) across 10 simulations comprising 16 sessions. **(E)** True and Viterbi-decoded states in a representative example session. **(F)** We assessed the Viterbi algorithm’s ability to identify state-transitions by quantifying – for each decoded transition – the distance to the nearest true transition point. The majority of transition-points were exactly identified (*M_distance_* = −0.03, *SD* = 6.45) and 84% of decoded transitions occurred within ±3 trial window of a true transition. Histogram shows frequency of decoded-vs-true transition differences.

**Figure S4.**
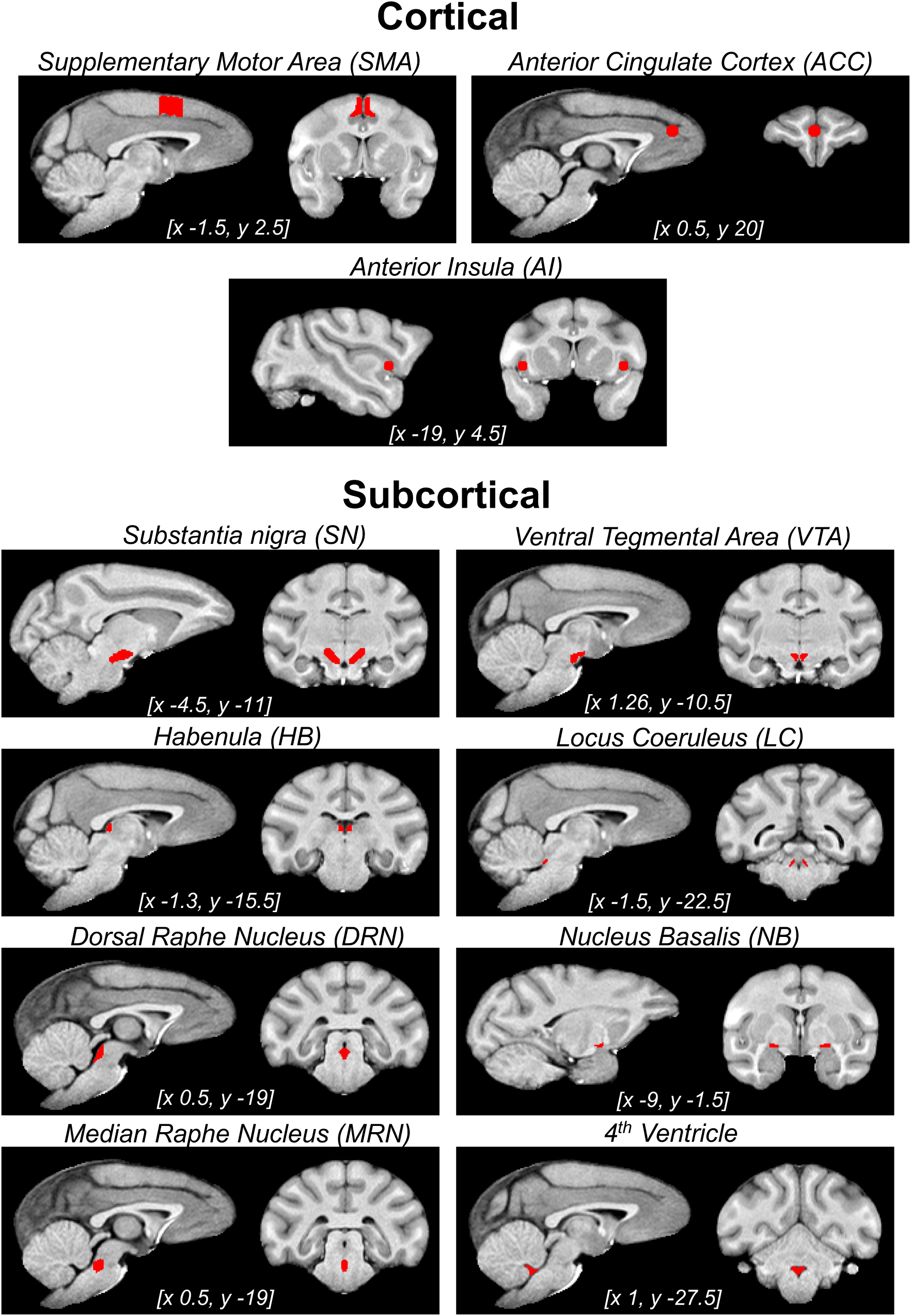
Cortical and subcortical regions of interest. An illustration of the ROIs implemented during fMRI analysis in CARET f99 macaque space. Subcortical and SMA ROIs consisted in anatomical masks that were drawn on a group structural template in CARET F99 macaque monkey space and then warped to individual structural and functional spaces by nonlinear transformation. These masks were constructed separately by two-different assessors based on the Rhesus Monkey Brain Atlas^72^ and then evaluated on for convergence across assessors. ACC and AI ROIs were defined as 3mm spheres centred on the peak of functionally relevant activation contrasts obtained in previous studies^19,50^. The medial raphe nucleus (MRN) and fourth ventricle were only used as control ROIs to investigate the focality of the DRN effect.

**Figure S5.**
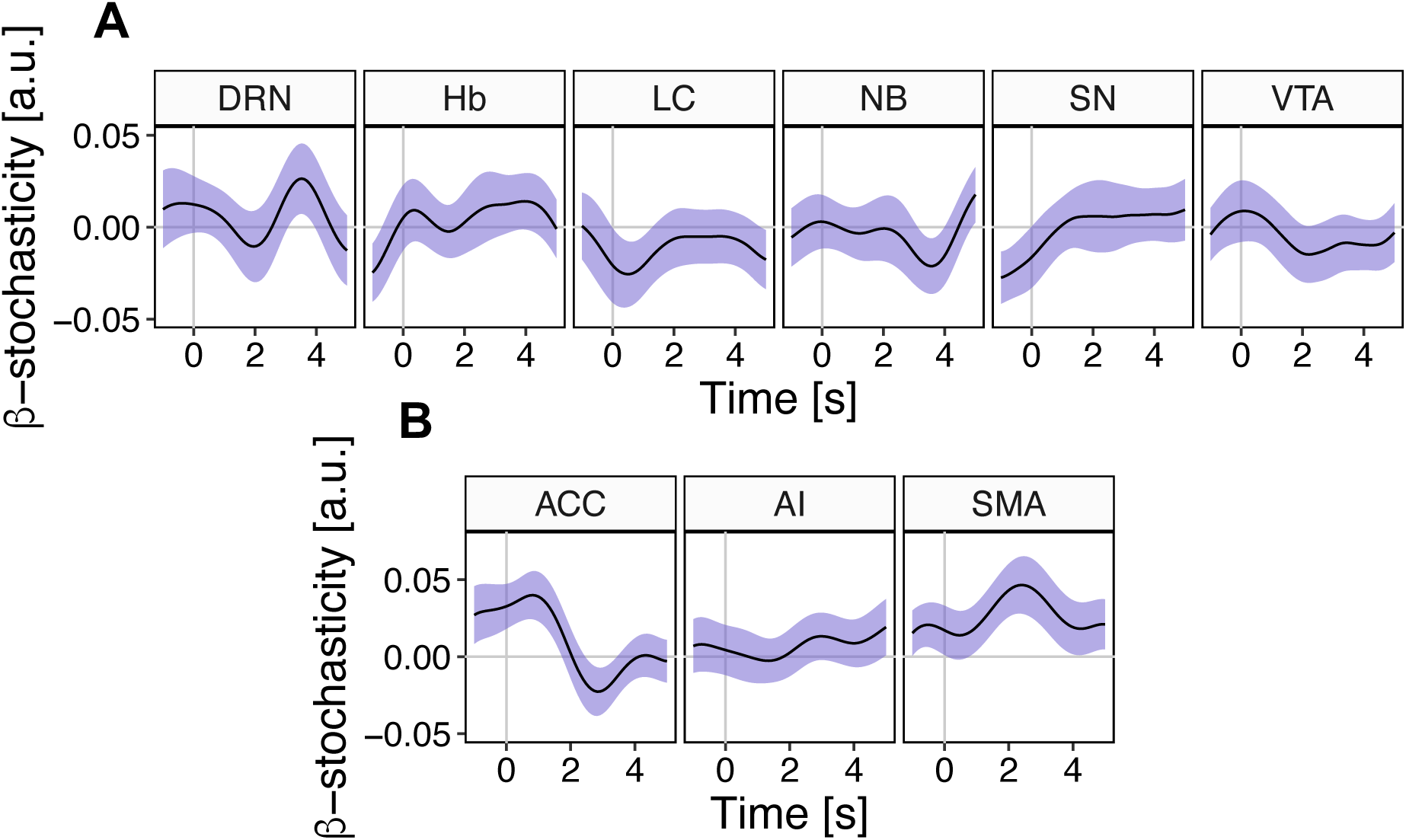
No evidence for reward-stochasticity representations in ROIs. The time-course and peak-regression-coefficients for the effect of reward-stochasticity on blood oxygen level dependent (BOLD) signal in subcortical **(A)** and cortical **(B)** ROIs. Epochs are time-locked to decision-making. Reward-stochasticity had a modest influence on behaviour but there was no evidence that it was represented in the brain activity of ROIs (GLM3.1).

**Figure S6.**
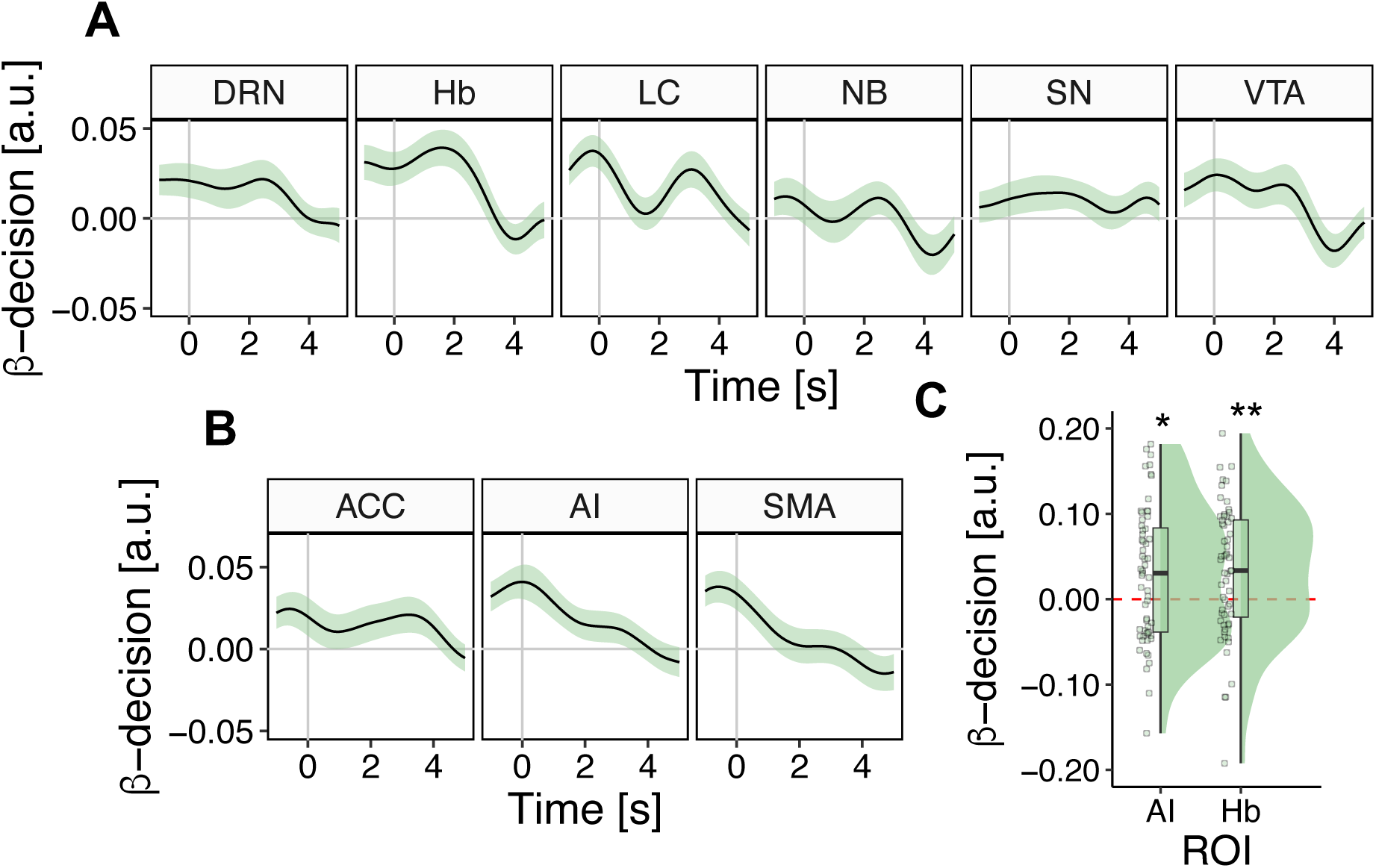
Habenula and Anterior Insula encode pursue/reject decisions. The time-course and peak-regression-coefficients of the relationship between pursuit decisions (pursue-vs-reject) and BOLD signal in subcortical **(A)** and cortical **(B)** ROIs. Epochs are time-locked to decision-making. **(C)** Pursuit decisions were represented in the BOLD activity of Hb (*t*_HB_(58) = 3.25, p = .012) subcortically and AI (*t*_AI_(58) = 2.78, p = .022) in the cortex. No other ROIs represented pursuit decisions.

**Figure S7.**
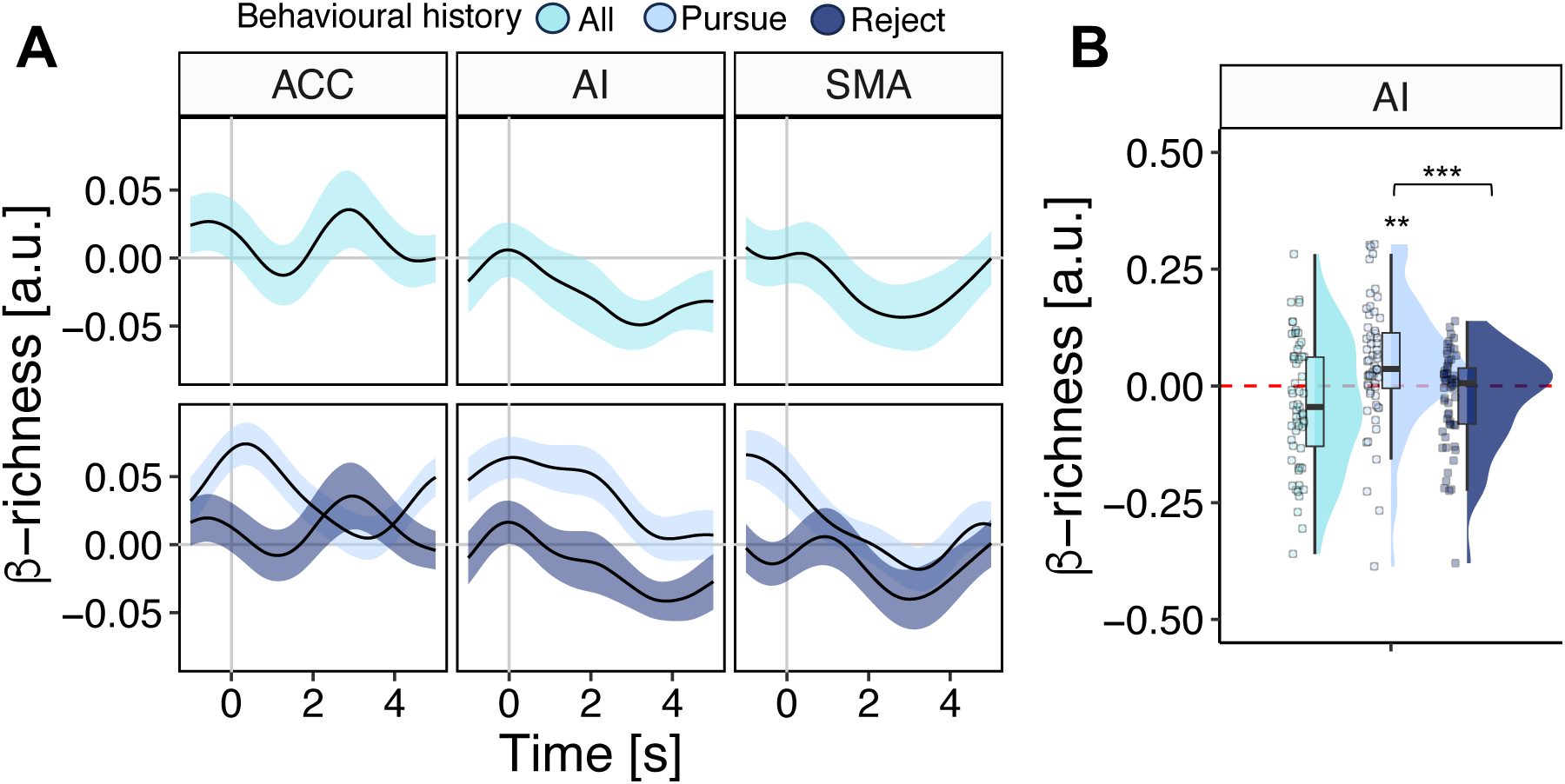
Anterior Insula represents the richness of the environment with the same pattern as subcortical ROIs. The time-course and peak-regression-coefficients of the relationship between richness of the environment and BOLD signal in cortical ROIs. All epochs are time-locked to decision-making. AI is the only cortical ROI to represent an animal’s environment as a function of pursue/reject behaviour (GLM3.2; *t_AI; rejected_*(58)=-1.94, *p*=.057; *t_AI; pursued_*(58)=3.02, *p*=.004; *t_AI; pursued-vs-rejected_*(58)=3.73, *p*<.001; Holm-Bonferroni correction not applied).

**Figure S8.**
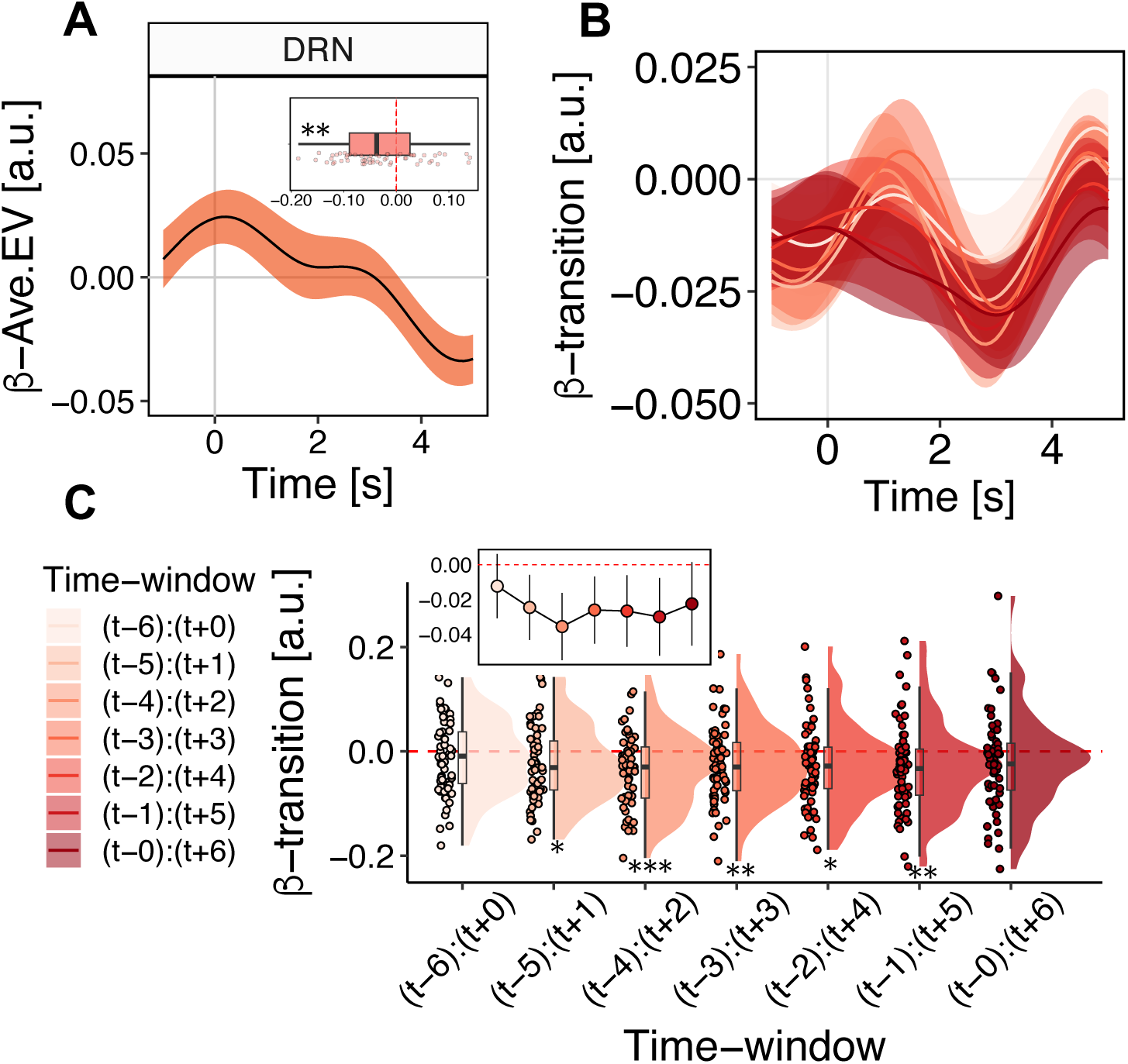
Further examination of motivation-state transitions in Dorsal Raphe Nucleus. **(A)** Our analysis of behaviour indicated that animals were more likely to occupy high motivation-states as the average expected-value of recently encountered rewards increased (fig. 2J). Given that BOLD activity in DRN represented transitions between motivation-states, we tested whether DRN also encoded the average expected-value of recent reward opportunities – the idea being that DRN might control changes in motivation-states specifically in relation to the distribution of available rewards. This was, indeed, the case – the average expected-value of the preceding five reward opportunities exerted a negative effect on BOLD activity in DRN (*t_EV_*(58) =-2.89, p=.005; GLM3.5). Inset panel shows distribution of peak regression coefficients. **(B&C)** We initially tested the effect of motivation-state transitions on brain activity in a symmetric 7 trial window centred on the transition-trial (i.e. the transition trial +/− 3 trials). This indicated that transitions were represented in the BOLD activity of DRN, as reported in the main text. We confirmed that this effect was not an artefact of our initial window selection by iterating the analysis over different 7-trial windows comprising transitions events. This showed that the effect was robust to window position (*t*_(t-5):(t+1)_(58)=-2.63, p=.011; *t*_(t-4):(t+2)_(58)=-3.69, p<.001; *t*_(t-3):(t+3)_(58)=-2.70, p=.009; *t*_(t-2):(t+4)_(58)=-2.60, p=.012; *t*_(t-1):(t+5)_(58)=-2.69, p=.009; *t*_(t-0):(t+6)_(58)=-1.88, p=.065; GLM3.6). Note that the windows featuring the least overlap with decoded motivation-transitions show non-significant effects (*t*_(t-6):(t+0)_(58)=-1.34, p=.180; *t*_(t-0):(t+6)_(58)=-1.88, p=.065). Inset panel in D shows mean ± 95% confidence interval of transition effect across time-windows.

**Figure S9.**
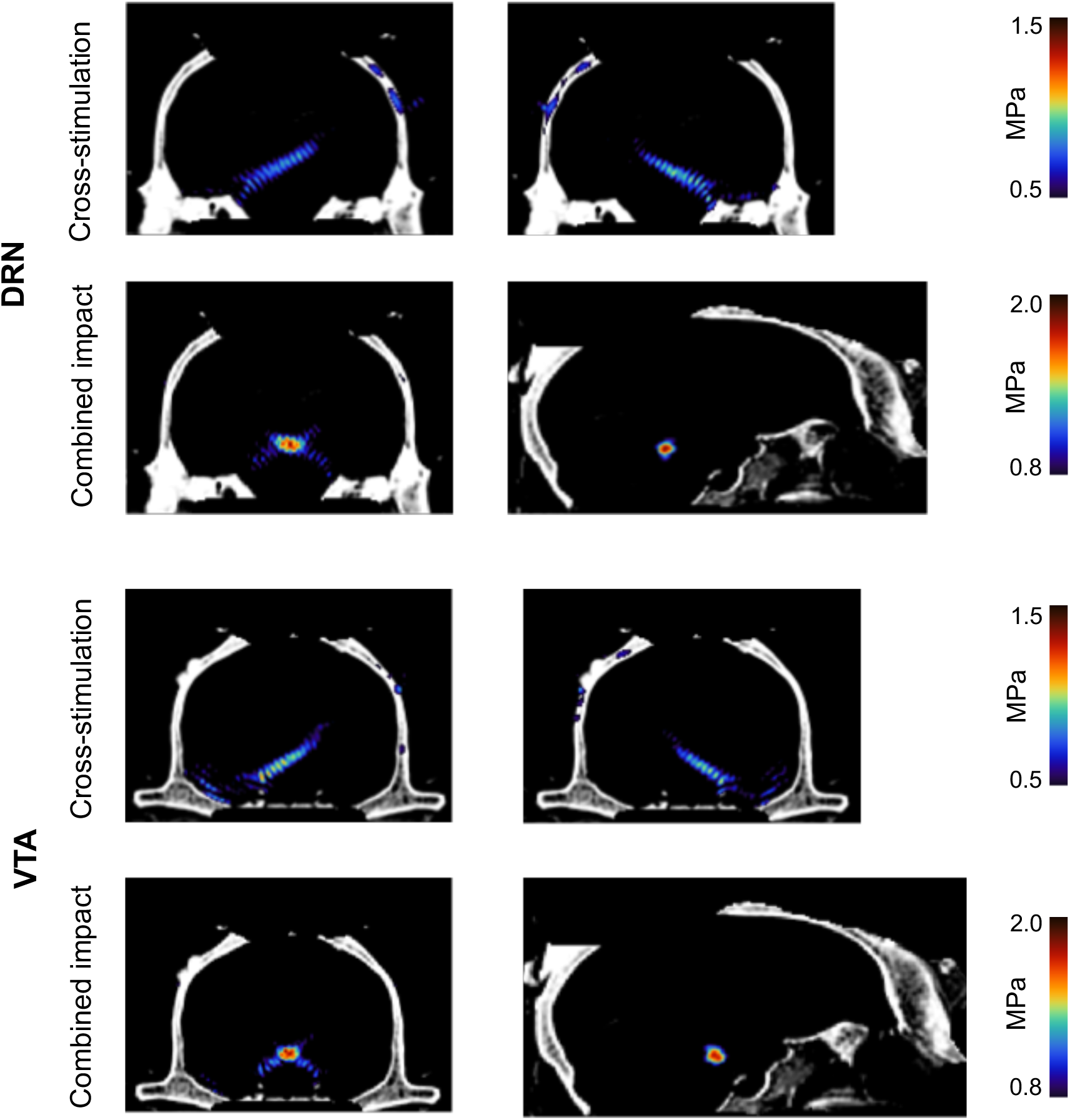
Simulation of the propagation of acoustic waves produced by the TUS protocol targeting DRN and VTA. Simulated focused ultrasound peak intensities and spatial distribution in the brain from successive left and right DRN / VTA stimulation. The skull was estimated from pseudo-CT images obtained from each monkey using a Black Bone MRI sequence ^73^. The maximum spatial-peak pulse-averaged intensity (Isppa) at the acoustic focus point was 24.4 W/cm^2^ (0.86 MPa) for the left DRN target, 21.5 W/cm^2^ (0.80 MPa) for the right DRN target, 30.3 W/cm^2^ (0.95 MPa) for the left VTA target and 19.9 W/cm^2^ (0.77 MPa) for the right VTA target. The combined impact is calculated as the sum of stimulation intensity map across the two consecutive stimulations delivered over the left and right hemisphere for the DRN and VTA targets. The simulation procedure is described in the Methods and previous studies^74^. The simulated data shown here is from monkey M2.

### Experimental model and subject details

The experiments were performed with four male rhesus macaques (*Macaca mulatta*). The sample size corresponded to those used in previous studies in which it had been possible to identify significant and reliable whole-brain functional magnetic resonance (fMRI) recording and transcranial ultrasound stimulation (TUS) across multiple testing sessions but with the minimum number of animals. The animals were 10-12 years of age and weighed 14.30–17.09 kg. They lived in group housing with a 12hr light dark cycle and were afforded access to water for 12–16 hr on testing days and free access on non-testing days. All procedures were conducted under a license issued by the UK Home Office in accordance with the Animal (Scientific Procedures) Act 1986 and the European Union guidelines (EU Directive 2010/63/EU).

### Behavioural training

All monkeys were fitted with magnetic resonance imaging (MRI) compatible cranial implants that facilitated head-fixation during testing and training (Rogue Research). Training was conducted in MRI compatible chairs designed to accommodate monkeys in a sphinx position and took place in custom-built environments that replicated an MRI scanner. Animals were trained on simplified versions of the task in which they did not need to wait go-cue and action-outcome delays and were not exposed to controlled changes in their reward environment. Go-cue and action-outcome delays were then introduced and carefully increased over the course of training until they were sufficient for an fMRI experiment (see below). Changes in the reward environment were introduced after animals were accustomed to the final delay timings. Testing for both the fMRI and transcranial ultrasound stimulation (TUS) experiments began when the proportion of pursued trials per session was stationary over consecutive days.

### Behavioural task

Animals performed a simple decision-making task involving sequential encounters with reward opportunities that appeared on a computer screen. Reward opportunities were presented in visual form via coloured boxes (size of the box: 8 x 26 cm) that were filled with dots. The colour of the stimulus indicated the number of juice drops that the opportunity was worth {red = 1; green = 2; blue = 3}. The number of dots comprising the stimulus indicated the probability that reward would be delivered if the opportunity was pursued. Reward probabilities ranged between (.05, 1), and dots indicated .05 linear increments of reward probability. The stimuli were designed so that dots gradually filled the box from the top downwards.

Opportunities first appeared in the centre of the screen before displacing either right or left (fig. 1A). The displacement of the opportunity-stimulus functioned as a go-cue which indicated that the opportunity was available for pursuit. Monkeys could pursue opportunities by manually responding to an infrared sensor corresponding to the opportunity’s on-screen location – for example, if the opportunity displaced right, they needed to touch a sensor with their right-hand, and vice versa if it displaced left. The go-cue was designed to temporally dissociate decisions about pursuing opportunities from the motoric processes which realised decisions in behaviour. Similarly, the randomised right/left displacement of the opportunity prevented monkeys from motor planning during the decision phase. The durations separating opportunity-onsets and go-cues were drawn from *go-cue ∼ uniform(3, 4)* and were optimised for a canonical macaque hemodynamic response function (HRF) of approximately 4s. Monkeys had 1s to pursue opportunities after the go-cue. After pursuing opportunities monkeys needed to wait action-outcome delays spanning *A-O-delay ∼ uniform(3, 4)* before receiving potential juice rewards accompanied by visual reward feedback. The next trial then began after an inter-trial interval spanning *ITI ∼ N(4, 1)*. If monkeys did not pursue an opportunity, they bypassed the action-outcome and reward-delivery durations and proceeded to the next trial. If monkeys pursued opportunities before the go-cue – i.e. prematurely – they needed to wait the remaining duration of the opportunity presentation, in addition to go-cue, action-outcome and reward-feedback durations. The opportunity which elicited the premature response was then repeated on the next trial.

Each session of the task comprised four blocks of 40–50 trials. Blocks were used to engender different reward environments by systematically controlling the reward-probability aspect of opportunities. Environments varied on two dimensions: (i) *richness* was defined by the mean of the reward-probability distribution, where μ_rich_ = 0.75 and μ_poor_ = 0.55, and (ii) stochasticity was defined by the width of reward-probability distributions, where predictable environments were generated from Gaussian distributions with σ_predictable_ = 0.05 and unpredictable environments from random-uniform distributions with ranges of 0.40 such that σ_unpredictable_= 0.13. Blocks were characterised along both richness and stochasticity dimensions, which yielded four distinct environment-types – rich-predictable, rich-stochastic, poor-predictable, and poor-stochastic. Each session featured one of each environment-type, and the order of environments was counterbalanced with respect to the richness dimension to avoid long periods of low-value offers (e.g., during consecutive poor environments) which were difficult for monkeys to perform. Environment-types were indicated by visual cues which bordered the screen.

The task was implemented in MATLAB v2019 (MathWorks) using Psychophysics Toolbox v3 ^75^ and presented on MRI-compatible screens (23in BOLD screen; Cambridge Research Systems) approximately 30cm away from the subjects. Juice rewards consisted in a solution of water, blackcurrant cordial and banana and each juice drop was approximately 1mL. Pupillometric data was obtained during fMRI sessions via an MR-compatible infrared EYElink 1000 eye-tracker device (SR Research Ltd.) recording pupil-diameter and gaze-direction along x and y planes at a 250-Hz sampling rate.

Pre-processing and analysis of pupillometric data was performed in *R* and entailed the following steps (adapted from ^76^). Samples reflecting eye blinks were identified with a detection algorithm native to the EYElink device. Artefacts were defined as consecutive samples with discrepancies of > 50 a.u. All samples within symmetric 25 sample (i.e., 0.1s) windows of artefacts or eye blinks were scrubbed and linearly interpolated, and interpolated time series were then low pass filtered with a 4Hz cut-off. These processes were performed for pupil-size, x-gaze-direction, and y-gaze-direction time series respectively, which were z-scored. We then regressed variance due to x-gaze-direction and y-gaze-direction from pupil-size via linear regression and performed subsequent analysis using the residuals.

To incorporate pupillometry in our behavioural analysis, we extracted pupil-size in specific epochs of the task, with a focus on periods after the presentation of reward opportunities in which animals were making decisions. In particular, we quantified – for each trial – mean pupil-size 0.5–1.0s after stimulus presentation. We selected this epoch to avoid pupillometric responses reflecting sudden luminance changes, as when a visual stimulus first appears.

### Behavioural analysis

Our initial analysis of behaviour characterised the influences on binomial pursuit/reject decisions. We accomplished this with mixed-effect binomial generalised linear models (GLMs) with subject-identity as a random variable. We quantified the richness of an animal’s reward environment via a moving-average with a retrospective five-trial window, and the stochasticity of its environment as the standard-deviation of that reward rate as follows:

> **Richness of environment**
>
> *richness_t_ = mean(reward-received_t–5_: reward-received_t–1_)*

> **Stochasticity of environment**
>
> *stochasticity_t_ =* sd(*richness*_t–5_: *richness*_t–1_)

We implemented the following model to assess the factors influencing pursuit/reject decisions:

> **GLM1.1**
>
> *logit(decision_t_) = β_o_ + β_1_reward-magnitude_t_ + β_2_reward-probability_t_ + β_3_environment-richness_t_ + β_4_environment-stochasticity_t_ + β_5_ (reward-magnitude_t_*reward-probability_t_) + β_6_ (trial-number_t_) + μ_0_ + ε*

where β_0-6_ are fixed-effects, μ_0_ is the by-subject random intercept and ε is an error term. The small sample size precluded fitting GLM1.1 with the full random-effects structure – that is, affording all predictors random effects by subject. We therefore fit an iterative series of models in which the key predictors of interest – *environment*-*richness* and *environment-stochasticity*– were individually afforded random effects by animal. All mixed-effects GLMs were performed in *R* and models were fit via maximum likelihood estimation as implemented in the *lme4* toolbox. We then fit the following GLMs in using the same procedure:

> **GLM1.2**
>
> *logit(decision_t_) = β_o_ + β_1_reward-magnitude_t_ + β_2_reward-probability_t_ + β_3_behavioural-history_t_ + β_4_ (reward-magnitude_t_*reward-probability_t_) + β_5_trial-number_t_ + μ_0_ + ε*

where β_0-5_ are fixed-effects, μ_0_ is the by-subject random intercept and ε is an error term.

> **GLM1.3**
>
> *logit(decision_t_) = β_o_ + β_1_reward-magnitude_t_ + β_2_reward-probability_t_+ β_3_behavioural-history_t_ + β_4_environment-richness_t_ + β_5_ (environment-richness_t_*behavioural-history_t_) + β_6_ (trial-number_t_) + μ_0_ + ε*

where β_0-6_ are fixed-effects, μ_0_ is the by-subject random intercept and ε is an error term.

### Behavioural modelling

We used an GLM-HMM to probe time-varying patterns in behaviour that might reflect internal motivation-states. A GLM-HMM is an extension of the GLM that consists of two parts. The HMM part of the model assumes that time series events arise from so-called hidden states, which produce observations according to state-dependent probability distributions ^26,27^. These states bear Markov relations to one another, such that the state at time t is determined strictly by the state at time t-1, and the respective probabilities of transition between states. A simple HMM of animal behaviour in this context, for example, might comprise states in which responses arise from binomial distributions with state-dependent probability parameters that reflect changes in an animal’s motivation (i.e., probability of pursuing) across time.

The GLM part of a GLM-HMM parameterises state-dependent probability distributions according to predictors^26^. The weights afforded to each predictor can vary from state-to-state, which enables the model to capture time-varying differences in an animal’s decision-making process. We implemented a GLM-HMM in which pursuit/reject decisions were parameterised by bias terms, in addition to predictors for the expectation-value of the specific opportunity animals were faced with on the current trials and binary environment cues indicating the experimentally manipulated richness and stochasticity of the reward environment. We were interested in internal states of motivation – that is, changes in an animal’s intrinsic propensity to pursue rewards. We therefore let only the bias parameter(s) change between states and constrained the weight on expectation-value and environment-cue parameters to be the same across states. Formally:

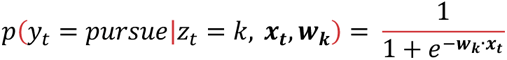

Where y ∈ {pursue, reject}^T^ is the animal’s decisions across *T* trials, *z_t_* is the HMM state at trial *t* and z_t_ ∈ {1,…, K} such that there are *K* states,, *x*_t_ ∈ ℝ^M^ is a matrix of *M* predictors in the GLM part of the model at time *t* and ***w***_k_ ∈ ℝ^M^ represents *M* GLM weights over predictors that are specific to state k. Transitions between the states unfold according to a transition matrix *A* ∈ ℝ^K x K^:

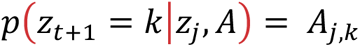

where transition matrix is stationary. The joint probability of behaviours and HMM states is given by:

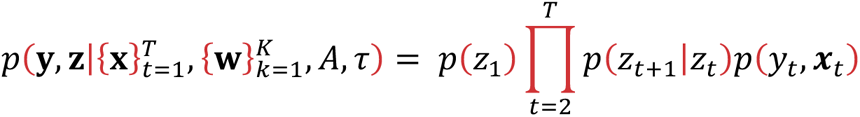

Where τ ∈ ℝ^K^ is the initial state distribution such that τ*_k_* = *p*(*z*_1_ = *k*). The likelihood function was fit to behavioural data via Markov Chain Monte Carlo optimisation implemented in STAN ^77^. For fMRI sessions, GLM-HMMs were fit to each individual animal’s behaviour. For TUS sessions, GLM-HMMs were fit for each stimulation condition.

We validated the GLM-HMM by confirming parameter recovery for diversely configured generative models (see fig. S2 & fig. S3). We fit an iterative series of models with *K* ∈ {1, 2, 3, 4, 5} states to behaviour. We evaluated these models in following ways. Firstly, we compared their scores on session-wise log-likelihood and AIC – a well-established and easily interpretable metric of model fit that penalises complexity. Secondly, we implemented a five-fold cross-validation protocol in which GLM–HMMs were iteratively fit to 80% of sessions for each animal and tested on the remaining 20% of sessions ^26^. We then calculated the log-likelihood of held-out sessions (see fig. S2). We further validated GLM-HMMs by generating simulated datasets from fitted models. We then inspected these simulations for key features and patterns of animal behaviour (fig. S2).

After performing model selection on GLM-HMMs, we decoded the maximum a posteriori sequence of HMM-states in each session via the Viterbi algorithm ^77^.

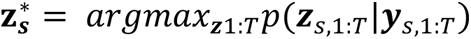

Decoding the HMM-states allowed us to quantify their impact on behaviour, which we did using the following mixed-effects GLMs with subject-identity as the grouping variable:

> **GLM2.1**
>
> *logit(decision_t_) = β_o_ + β_1_reward-magnitude_t_ + β_2_reward-probability_t_ + β_3_motivation-level_t_ + β_4_ reward-magnitude_t_*reward-probability_t_ + β_5_trial-number_t_ μ_0_ + μ_1_motivation-level_t_* + *ε*

where β_0-5_ are fixed-effects, μ_0-1_ are by-subject random effects for intercepts and motivation-level, respectively, and ε is an error term. Motivation-level ∈ {0,1} is a binary variable reflecting the motivation-state (low vs high) occupied on trial *t*. We identified motivation-state transitions by searching for cases where decoded motivation-states were different across consecutive trials (i.e., *motivation-level_t_ ≠ motivation-level_t-1_)*. We compared behaviour before-vs-after motivation-state transitions with the following mixed-effects GLM:

> **GLM2.2**
>
> *logit(decision_t_) = β_o_ + β_1_reward-magnitude_t_ + β_2_reward-probability_t_ + β_3_before-vs-after-transition_t_ + β_4_ reward-magnitude_t_*reward-probability_t_ + β_5_trial-number_t_ μ_0_ + μ_1_motivation-level_t_* + *ε*

where β_0-5_ are fixed-effects, μ_0-1_ are by-subject random effects for intercepts and time-from-transition, respectively, and ε is an error term. *Before-vs-after-transition* was a binary variable that grouped trials into two categories: (i) ‘before-transition’ trials, defined as the five trials preceding a motivation-state transition, and (ii) ‘after-transition’ trials, defined as the five-trials in the immediate aftermath of a transition. We performed GLM2.2 separately for low-to-high and high-to-low motivation-state transitions. We then characterised the GLM-HMM’s convergent validity by comparing decoded states with variables that should change with motivation: (i) reaction-time, and (ii) pupil-size. To do this, we implemented the following mixed-effects GLMs:

> **GLM2.3**
>
> *Pupil-size_t_ = β_o_ + β_1_reward-magnitude+ β_2_reward-probability+ β_3_HMM-state + β_4_trial-number + μ_1_HMM-state + ε*

> **GLM2.4**
>
> *log(Reaction-time _t_) = β_o_ + β_1_reward-magnitude+ β_2_reward-probability+ β_3_HMM-state + β_4_trial-number + μ_1_HMM-state + ε*

Where β_0-4_ are fixed-effects and μ_1_ are random-effects within each subject. We did not implement random intercepts (i.e. μ_0_) for either model because *pupil-size* and *reaction-time* were z-scored within-subjects during pre-processing. This meant that the central tendency of within-subject distributions for *reaction-time* and *pupil-size* was the same value (i.e., 0), and it was not necessary to add terms capturing inter-subject differences.

We used changes in decoded HMM-states to examine the temporal dynamics of motivation states. We were particularly interested in links between motivation-states and aspects of an animal’s external environment, like the recent history of reward opportunities it had been presented with. We quantified the latter by calculating a moving average of the expectation-value of reward opportunities received in the previous five trials.

> **Availability of rewards (Ave. EV)**
>
> *Ave. EV_t_ = mean(EV_t–5_: EV_–1_)*

We used Ave. EV to predict motivation-states via the following mixed-effects Binomial GLM:

> **GLM2.5**
>
> *logit(motivation-state_t_) ∈ {0,1} = β_o_ + β_1_EV + β_2_Ave.EV + β_3_trial-number + μ_0_ + μ_1_Ave.EV + ε*

Where β_0-3_ are fixed-effects and μ_0-1_ are random-effects within each subject. We used the EV – the expected value of the current offer on a given trial – to control for the effects of the current offer on an animal’s motivation-state. We used EV rather than the separate reward-magnitude and reward-probability dimensions of offers because Ave. EV and reward-probability were meaningfully correlated (r = 0.49). This correlation was an inherent feature of the task because reward environments were engendered by manipulating the distribution of reward-probabilities over time. This created autocorrelations in reward-probabilities over time and, therefore, correlations between reward-probability and retrospective summary statistics of task events like the Ave.EV. Scaling reward-probabilities by reward-magnitude as in the expectation-value formula dampened this correlation (*r* = 0.23), meaning that EV and Ave. EV could be part of the same regression without collinearity.

### Acquisition, reconstruction and pre-processing of MRI data

During collection of fMRI data monkeys were head-fixed in a sphinx posture within an MRI compatible chair (Rogue Research). MR images were acquired with a horizontal bore clinical 3T scanner with a 15-channel non-human primate (NHP)-specific receive coil (RAPID Biomedical). Structural images were acquired during a previous experiment^50^. Functional images were acquired via the CMMR multiband gradient-echo T2* echo planar imaging (EPI) sequence designed specifically to achieve high signal-to-noise (SNR) in subcortical structures^78,79^. This was characterised by 1.25mm isotropic voxels with a repetition time (TR) of 1282ms, echo time (TE) of 25.40ms, multiband acceleration factor MB=2, in-plane acceleration factor R=2, and flip angle of 63°.

Offline reconstruction of the raw functional data was performed following the dynamic off-resonance correction method developed by Shahdloo et al^80^. In summary, standard Nyquist ghost correction and dynamic zeroth-order B0 correction were applied first. Then, the EPI reference navigator data acquired at every time-point was compared to navigator data from single-band references to estimate first-order dynamic off-resonance perturbations arising from the awake animal’s body movements. Finally, the off-resonance estimates were used to correct the raw data prior to reconstruction.

Pre-processing of MR images was performed with a combination of FMRIB’s software library, Advanced Normalization Tools, the Human Connectome Project Workbench and Magnetic Resonance Comparative Anatomy Toolbox^81^. Although monkeys were head-fixed during MRI acquisition, incidental limb and body movements caused time-varying distortions in the B_0_ magnetic field and therefore non-linear motion artefacts along the phase encoding direction. To account for this, a low-noise EPI volume was identified for each session and then implemented as a reference to which other volumes were non-linearly registered slice-by-slice along the phase-encode direction. Aligned and distortion-corrected EPIs were then registered non-linearly first to monkey-specific high-resolution images, and then to a group template in CARET f99 macaque space. Further details of the group template construction are described elsewhere^51^. Finally, the functional images were temporally filtered (high-pass temporal filtering, 3-dB cutoff of 100s) and spatially smoothed (Gaussian spatial smoothing, full-width half maximum of 2.5mm).

Three measures were used to detect artefacts in the data: a) For each slice in each volume the linear transform (in the y-plane) from that slice to the corresponding slice in the mean reference image; b) The normalized correlation between that slice and the corresponding slice in the mean reference image; c) For each volume, the correlation between that volume (mean-filtered across z-slices) and the mean reference image after correction. Volumes were removed when they exceeded 2.5 SDs above the median of each measure. The threshold was chosen to keep the number of censored volumes less than 10% of the total volumes. We also added 13 PCA components describing, for each volume, the warping from that volume to the mean reference image when correcting motion artefacts (i.e., they capture signal variability associated with motion induced distortion artefacts), as parametric regressors of non-interest that were not convolved in our general linear models.

### fMRI data analysis

We focused our analysis of fMRI data on circumscribed *a priori* regions of interest (ROIs) comprising dorsal raphe nucleus (DRN), ventral tegmental area (VTA), substantia nigra (SN), nucleus basalis (NB), locus coeruleus (LC) and habenula (Hb) in the subcortex, and anterior cingulate cortex (ACC), anterior insula (AI) and supplementary motor area (SMA) in the cortex. Subcortical and SMA ROIs consisted in anatomical masks that were drawn on a group structural template in CARET F99 macaque monkey space and then warped to individual structural and functional spaces by nonlinear transformation. These masks were constructed separately by two-different assessors based on the Rhesus Monkey Brain Atlas^72^ and then evaluated on for convergence across assessors. ACC and AI ROIs were defined as 3mm spheres centred on the peak of functionally relevant activation contrasts obtained in previous studies^19,50^.

We extracted the filtered time series of BOLD signal from each ROI. The extracted signals were then averaged, normalised and up-sampled by a factor of 15^19,50,51^. The upsampled data was then epoched to 6s time windows spanning 1s before to 5s after the appearance of the reward-opportunity stimulus on each trial. We then examined the relationship between behaviour and brain activity with ordinary least squares (OLS) GLMs performed at each timepoint in each epoch.

Inferential statistics for time-course GLMs were performed using a leave-one-out cross-validation (LOOCV) procedure designed to estimate the peak regression-coefficient in each session without selection bias^19,50,51^. For each session *s* (*N* = 59), we determined the timepoint *t* at which the largest absolute-value regression coefficient occurred in the remaining *N-1* (i.e., 58) sessions. We restricted our search for *t* to a 4s window from 1s after to 5s after the decision-making time – that is, a four second window centred on the mean macaque HRF. We then calculated the regression coefficient in session *s* at time *t*. We repeated this iteratively for each session, which yielded a series of 59 regression coefficients. We performed significance testing on regression coefficients with single-sample t-tests. To control for multiple comparisons, we applied the Bonferroni-Holm correction for any analysis performed on more than one ROI. For each GLM, we repeated this for each regressor in each ROI. All time course analysis was conducted in MATLAB (MathWorks) using custom analysis scripts.

We implemented the following series of GLMs:

> **GLM3.1**
>
> *BOLD_ROI_ = β_o_ + β_1_reward-magnitude+ β_2_reward-probability + β_3_environment-richness + β_4_environment-stochasticity + β_5_behavioural-history + β_6_environment-richness*behavioural-history + β_7_environment-stochasticity*behavioural-history + β_8_pupil-size + β_9_trial-number*

Where *BOLD_ROI_* indicates a t-by-s matrix containing time-series data for a given ROI (where t is trial, and s is time-sample). Pupil-size and trial-number were added as confound regressors to control for the effects of arousal and fatigue, respectively.

> **GLM3.2**
>
> *BOLD_ROI_ = β_o_ + β_1_reward-magnitude+ β_2_reward-probability + β_3_environment-richness + β_4_environment-stochasticity + β_5_pupil-size + β_6_trial-number*
>
> GLM3.2 was performed separately for subsets of trials in which an animals pursued and rejected the previous opportunity, respectively.

> **GLM3.3**
>
> *BOLD_ROI_ = β_o_ + β_1_decision + β_2_pupil-size + β_3_trial-number*
>
> Where *decision* is the pursue vs. reject decision made on a given trial.

> **GLM3.4**
>
> *BOLD_ROI_ = β_o_ + β_1_transition-period + β_2_ motivation-state-identity + β_3_decision + β_4_pupil-size + β_5_trial-number*

Where *transition-period* is a dummy-coded variable covering symmetric 7-trials windows (*t-3:t+3*) around decoded motivation-state transitions and *motivation-state-identity* is the decoded motivation-state (high vs. low) for a given trial. GLM3.4 was subsequently performed with 7-trial transition-periods aligned to from (t-6:t+0) to (t-0:t+6) with respect to decoded transitions to compare the timing of transition-related signals (see fig.3 and supplementary figure S8). It was also performed separately for low-to-high and high-to-low transition events (see fig. 3).

We tested whether DRN coded the value of recently available rewards (Ave. EV) with the following GLM:

> **GLM3.5**
>
> *BOLD_DRN_ = β_o_ + β_1_EV + β_2_Ave. EV + β_3_pupil-size + β_4_trial-number*

GLM3.5 was performed on data from DRN alone, as it was designed to test a specific hypothesis arising from the combined neural and behavioural data. We used EV to control for the effects of the currently available offer instead of separate reward-magnitude and reward-probability regressors as in previous GLMs of BOLD signal. This is due to collinearity between Ave. EV and reward-probability regressors (see GLM2.5 for further explanation).

We probed changes in connectivity as a function of richness of the environment using the following PPI-GLM.

> **GLM3.6**
>
> *BOLD_ROI_ = β_o_ + β_1_BOLD_seed_ + β_2_PPI + β_3_environment-richness + β_4_reward-magnitude + β_5_reward-probability + β_6_environment-stochasticity + β_7_pupil-size + β_8_trial-number*

Where *BOLD_seed_ is* a t-by-s matrix containing time-series data for seed regions in PPI analysis, and *PPI* is the interaction between *BOLD_seed_* and *environment-richness* regressors. GLM 3.6 was performed separately for subsets of trials in which an animals pursued and rejected the previous opportunity, respectively. Analogously, we tested differences in functional connectivity with respect to (i) motivation-state level, and (ii) motivation-state transitions with the following PPI:

> **GLM3.7**
>
> *BOLD_ROI_ = β_o_ + β_1_BOLD_seed_ + β_2_PPI + β_3_transition-period + β_4_motivation-state-identity + β_5_pupil-size + β_6_trial-number*

Where *BOLD_seed_ is* a t-by-s matrix containing time-series data for seed regions in PPI analysis. The analysis was performed twice – once where *PPI* was the interaction between *BOLD_seed_* and *transition-period* regressors, and once where *PPI* was the interaction between *BOLD_seed_* and *motivation-state-identity* regressors. Finally, we tested differences in functional connectivity with respect to pursue/reject decisions with the following PPI:

> **GLM3.8**
>
> *BOLD_ROI_ = β_o_ + β_1_BOLD_seed_ + β_2_PPI + β_3_decision + β_4_pupil-size + β_5_trial-number*

Where *BOLD_seed_ is* a t-by-s matrix containing time-series data for seed regions in PPI analysis, and *PPI* is the interaction between *BOLD_seed_* and *decision* regressors.

### Transcranial ultrasound stimulation (TUS)

TUS was performed using a four-element annular array transducer (NeuroFUS CTX-500, 64mm active diameter, Brainbox Ltd, Cardiff, UK) combined with a programmable amplifier (Sonic Concept Inc’s Transducer Power Output System, TPO-105, Brainbox Ltd, Cardiff, UK). The transducer was paired with a transparent coupling cone filled with degassed water and sealed with a latex membrane. The water was degassed for 4-5 hr before each stimulation session and was replaced after each session. The resonance frequency of the ultrasonic wave was set to 500kHz. The stimulation protocol was based on previously established protocols in macaques (*9*, *18*, *44*). We used the following protocol: duty cycle 30%; pulse length 30ms; pulse repetition interval 100ms; total stimulation duration 30s. The pressure field from the transducer was measured in a water tank with a 75 µm diameter PVDF needle hydrophone (Precision Acoustics, Dorset UK) which had been calibrated at 500 kHz by the National Physical Laboratory (Teddington UK). The free-field spatial-peak pulse-average intensity (Isppa) at 60mm focal depth was 120 W/cm2, which was consistent with the output given by the transducer manufacturer.

At the beginning of each stimulation session the animal’s skull was shaved and a conductive gel (SignaGel Electrode; Parker Laboratories Inc.) was applied to the skin. The water-filled coupling cone and the gel was used to ensure ultrasonic coupling between the transducer and the animal’s head. Next, the ultrasound transducer / coupling cone was placed on the skull and a Brainsight Neuronavigation System (Rogue Research, Montreal, CA) was used to position the transducer so that the focal spot would be cantered on the targeted brain region. There were four stimulation conditions: 1) DRN (target of interest); 2) VTA (sub-cortical active control condition); 2) superior-temporal-sulcus (STS; cortical active control condition); 4) sham (passive control condition). DRN and VTA targets were approximately 50mm, and the STS 35mm, from the surface of the transducer (the exact focal distance depended on the subject). All targets were sonicated bilaterally for 60s in total, with 30s of stimulation applied to a target from each hemisphere. Sonication of the midline targets (DRN and VTA) from one hemisphere was immediately followed by sonication of the same target from the contralateral hemisphere (cross-beam stimulation; ^50^). Sonication of the STS in one hemisphere was immediately followed by sonication of a homologous target in the contralateral hemisphere. Hemispheres were sonicated in a pseudo-random order. After stimulation, monkeys were immediately moved to a testing room for behavioural data collection. The sham condition completely matched a typical stimulation session (setting, stimulation procedure, neuro-navigation, targeting, transducer preparation and timing of its bilateral application to the shaved skin on the head of the animal) except that sonication was not triggered. During the sham session the montage was pseudo-randomly positioned to target DRN, VTA or STS. Each stimulation condition was repeated five times, on separate days, and the order of the stimulation sessions was pseudo-randomized for each animal. The stimulation was always performed at the same time of the day and there was always a 24-hour gap between each session, regardless of it being a real or sham stimulation session.

### Acoustic modelling

We simulated the propagation of acoustic waves produced by the TUS protocol as described by Yaakub et al., 2023. In brief, we used k-wave – a k-space pseudospectral solver ^83^ – and kArray tools to obtain estimates for the pressure amplitude, peak intensity and spatial distribution of TUS at stead-state. First, we simulated the acoustic wave propagation in water (free field) to characterise ultrasound beam for a target intensity of 120 W/cm^2^ at 50mm focal depth (focal depth of VTA and DRN from transducer). Next, we performed the simulation for DRN and VTA targets in the skull. The skull was estimated from pseudo-CT images obtained from each monkey using a Black Bone MRI sequence ^73^. The skull was obtained by thresholding the pseudo-CT images at 1400-2100 Hounsfield Units (HU). A linear relationship between the pseudo-CT Hounsfield Units and the sound speed, as well as the density, and absorption coefficient was assumed as described elsewhere ^84,85^. The simulation grid size was set to the size of the T1-weighted MRI with a grid spacing of 0.5 mm which results in approximately 6 points per wavelength in water and tissue and up to 12.4 points per wavelength in bone.

### Resting-state imaging data acquisition, pre-processing, and analysis

We further validated the TUS protocol by examining its effect on resting state coactivation patterns between VTA/DRN and key interconnected cortical and subcortical regions. Awake resting-state fMRI (rs-fMRI) data was acquired for all four monkeys (same animals as in Experiments 1 and 2) pre-vs-post DRN-TUS and VTA-TUS, respectively. Pre-processing and analysis of rs-fMRI data has been described elsewhere ^25,82^.

We characterised the effects of TUS on the coactivation of DRN/VTA and ROIs by comparing rs-fMRI data collected before DRN/VTA TUS with the rs-fMRI data collected immediately after DRN/VTA TUS. Pre-stimulation and post-stimulation rs-fMRI were collected on the same day. The impact of DRN/VTA TUS on coactivation patterns was quantified with seed-based connectivity analyses, which involved calculating a series of pairwise linear correlations in BOLD activity between a seed region (DRN or VTA) and the remaining ROIs. The resulting pre-stimulation connectivity fingerprints for DRN-TUS and VTA-TUS were then contrasted with post-stimulation connectivity fingerprints before DRN-TUS and VTA-TUS (see fig. 4C).

### TUS data analysis

We characterised the behavioural effect of TUS by examining its two-way interactions with key predictors of decision-making in a series of Binomial GLMs. In these GLMs, the effect for TUS-condition was constructed to compare each individual active stimulation condition (STS, VTA, DRN) to sham-TUS as a reference. On some occasions, we compared DRN-TUS to one of the other active stimulation conditions – for example, DRN-TUS vs STS-TUS. We did this by performing GLMs on subsets of data that included only the TUS-conditions of interest. These analyses are specifically noted in the main text.

We tested the following to examine how TUS modulates the richness of the environment, and behavioural history effects, respectively.

> **GLM4.1**
>
> *logit(decision_t_) = β_o_ + β_1_reward-magnitude + β_2_reward-probability + β_3_environment-richness + β_4_environment-stochasticity + β_5_ TUS-condition + β_6_ (reward-magnitude*reward-probability) + β_7_ (environment-richness*TUS-condition) + β_8_ (trial-number) + μ_0_ + ε*

Where β_0-8_ are fixed-effects, μ_0_ is the by-subject random intercept and ε is an error term.

> **GLM4.2**
>
> *logit(decision_t_) = β_o_ + β_1_reward-magnitude + β_2_reward-probability + β_3_behavioural-history + β_4_ TUS-condition + β_5_ (reward-magnitude*reward-probability) + β_6_ (behavioural-history*TUS-condition) + β_7_ (trial-number) + μ_0_ + ε*

Where β_0-7_ are fixed-effects, μ_0_ is the by-subject random intercept and ε is an error term. The paucity of data in each TUS condition for each subject prevented us from fitting models with more complex random-effects structures for these models. We examined the influence of TUS on the frequency of transitions between motivation-states with the following GLM:

> **GLM4.3**
>
> *logit(state-transition_t_) = β_o_ + β_1_reward-magnitude + β_2_reward-probability+ β_3_TUS-condition + β_4_reward-magnitude*reward-probability + β_5_trial-number + μ_0_ + ε*

Where, state-transition ∈ {0,1}, β_0-5_ are fixed-effects, μ_0_ is the by-subject random intercept and ε is an error term. We performed a similar GLM to examine the influence of TUS on high-to-low transitions specifically:

> **GLM4.4**
>
> *logit(state-decrease_t_) = β_o_ + β_1_reward-magnitude + β_2_reward-probability+ β_3_TUS-condition + β_4_reward-magnitude*reward-probability + β_5_trial-number + μ_0_ + ε*

Where, state-decrease ∈ {0,1}, β_0-5_ are fixed-effects, μ_0_ is the by-subject random intercept and ε is an error term. We analysed TUS’s influence relationship between motivation-states and the availability of rewards (Ave. EV) with the following GLM:

> **GLM4.5**
>
> *logit(motivation-state_t_) = β_o_ + β_1_EV + β_2_Ave.EV + β_3_TUS-condition + β_4_TUS-condition*Ave.EV + β_5_trial-number + μ_0_ + ε*

Where, motivation-state ∈ {0,1} and the high-motivation=1 (i.e., positive coefficients correspond to increases in the likelihood of occupying the high-motivation state), β_0-5_ are fixed-effects, μ_0_ is the by-subject random intercept and ε is an error term. We performed GLM4.5 separately on behavioural data mean-split according to Ave.EV to determine whether DRN-TUS principally affected the influence between low-value environments and low-motivation states.

## Notes

### Competing Interest Statement

The authors have declared no competing interest.

